# Heat stress leads to rapid lipid remodelling and transcriptional adaptations in *Nicotiana tabacum* pollen tubes

**DOI:** 10.1101/2021.11.18.469149

**Authors:** Hannah Elisa Krawczyk, Alexander Helmut Rotsch, Cornelia Herrfurth, Patricia Scholz, Orr Shomroni, Gabriela Salinas-Riester, Ivo Feussner, Till Ischebeck

## Abstract

After reaching the stigma, pollen grains germinate and form a pollen tube that transports the sperm cells to the ovule. Due to selection pressure between pollen tubes, they likely evolved mechanisms to quickly adapt to temperature changes to sustain an elongation at the highest possible rate. We investigated these adaptions in *Nicotiana tabacum* pollen tubes grown *in vitro* under 22 °C and 37 °C by a multi-omic approach including lipidomic, metabolomic and transcriptomic analysis. Both glycerophospholipids and galactoglycerolipids increased in saturated acyl chains under heat stress while triacylglycerols changed less in respect to desaturation but showed higher levels. Free sterol composition was altered, and sterol ester levels decreased. The levels of sterylglycosides and several sphingolipid classes and species were augmented. Most amino acids increased during heat stress, including the non-codogenic amino acids γ-amino butyrate and pipecolate. Furthermore, the sugars sedoheptulose and sucrose showed higher levels. Also the transcriptome underwent pronounced changes with 1,570 of 24,013 genes being differentially up- and 813 being downregulated. Transcripts coding for heat shock proteins and many transcriptional regulators were most strongly upregulated, but also transcripts that have so far not been linked to heat stress. Transcripts involved in triacylglycerol synthesis were increased, while the modulation of acyl chain desaturation seemed not to be transcriptionally controlled indicating other means of regulation.

**One sentence summary:** *Nicotiana tabacum* pollen tubes react to heat stress with metabolomic adaptations, transcriptional adjustments, and a rapid and reversible lipid remodelling.

## Introduction

Around 80 % of the more than 350,000 known plant species are flowering plants (angiosperms) that mainly rely on sexual reproduction to pass on their genes to the next generation (Christenhusz and Byng, 2016). Sexual reproduction is also the prerequisite for seed and fruit set in crop plants. In most angiosperms, the male gametophyte – the pollen containing the sperm cells or the generative cell that later divides into the two sperm cells – double-fertilises the female gametophyte. To this end, the pollen has to land on the stigma of a pistil and adhere, rehydrate and germinate there. Since seed plant sperm cells are immotile, they have to be delivered through the female reproductive tissues to the ovule by tip-growing pollen tubes that are formed by the vegetative cells of the pollen (Sprunck, 2020). In some species, pollen tubes have to grow several centimetres to reach the egg-cell containing ovule. Maize pollen tubes, for example, can grow up to 50 cm in length with a speed of more than 1 cm per hour (Mascarenshas, 1993). Especially for these very long pollen tubes it is unlikely that enough lipids for the elongation are stored in the pollen grain, but the tubes rather depend on high *de novo* synthesis for their growth (Ischebeck, 2016). This assumption is supported by the fact that proteins involved in fatty acid synthesis are five to ten times more abundant in maturing pollen and growing pollen tubes of tobacco than in leaves or roots (Ischebeck *et al.*, 2014; Ischebeck, 2016).

The process of sexual reproduction in plants is very sensitive to abiotic stresses like heat stress (HS) or cold stress. Male gametophytes are especially sensitive to HS during all stages of their life and are generally more susceptible than female tissues (Hedhly, 2011; Santiago and Sharkey, 2019). One effect of HS on developing pollen is early tapetum degradation likely due to accumulation of reactive oxygen species (ROS) and the consequent disruption of concerted ROS signals (Zhao *et al.*, 2018). Furthermore, anthers can fail to release pollen grains, possibly due to heat-induced changes in cell wall composition, sucrose transport and water movement (Santiago and Sharkey, 2019). Also, the developing male gametophyte itself can encounter problems at different stages of development (Müller and Rieu, 2016; Raja *et al.*, 2019; Santiago and Sharkey, 2019).

The effects of HS on pollen germination and pollen tube growth are less well studied than those on pollen development but have also been addressed in several studies. It was shown that tomato pollen germination is negatively affected by HS in *in vitro* experiments (germination and pollen tube growth in medium outside of female tissues); an effect also attributed to elevated ROS levels (Luria *et al.*, 2019). Various sorghum genotypes show *in vitro* pollen germination reductions of 2-95 % under different heat stresses (Sunoj *et al.*, 2017). *In vivo* experiments with pollen from rice or the pine *Pinus edulis* gave similar results (Flores-Rentería *et al.*, 2018; Shi *et al.*, 2018).

Inhibited pollen tube growth has been observed as well. Experiments on *in vivo* grown cotton pollen tubes attribute reduced pollen tube growth to HS-induced reduction of soluble carbohydrate content in the pistil (Snider, Oosterhuis and Kawakami, 2011; Snider, Oosterhuis, Loka, *et al.*, 2011). Heat-induced pollen tube growth inhibition in rice was described to be likely due to altered auxin homeostasis within the pistil (Zhang *et al.*, 2018). These experiments highlight the importance of biochemical interactions between pollen tubes and the surrounding pistil tissue during stress. However, pollen tube growth inhibition is also observed *in vitro*.

Experiments with tomato pollen showed maximal germination rates at 15 °C and maximal pollen tube lengths at 25 °C, higher (or lower) temperatures were inhibitory (Karapanos *et al.*, 2010). Another *in vitro* study in tomato showed that high temperature-induced ROS inhibit pollen tube growth and that elevated flavonol levels can counteract the heat-induced ROS imbalance (Muhlemann, Younts and Muday, 2018). Also, *in vitro* pollen tube growth of cotton, rice and Arabidopsis is inhibited by high temperatures (Boavida and McCormick, 2007; Song *et al.*, 2015; Coast *et al.*, 2016), indicating that growth inhibitions are not solely due to altered crosstalk with the female tissue, but also due to effects in the pollen tube itself.

A challenge all organs of plants have to meet during elevated temperatures, is maintaining membrane fluidity and integrity; this holds true for the plasma membrane as well as for intracellular membranes (Niu and Xiang, 2018). Among the challenges plant membranes have to meet under elevated temperatures are the prevention of bilayer disintegration (because of membrane hyperfluidity under high temperatures) and peroxidation of unsaturated fatty acids by ROS (Higashi and Saito, 2019).

The effects of long- and short-term HS on the leaf lipidome of different species has been reviewed recently (Higashi and Saito, 2019; Lu *et al*., 2020): Levels of glycerophospholipids containing saturated or mono-unsaturated acyl chains increase; while most glycerophospholipid species with polyunsaturated acyl chains decrease. At the same time, triacylglycerol (TG) levels increase.

*Nicotiana tabacum* used in this study is a versatile model organism that is also very suited to study pollen tube growth. Here we show that tobacco pollen tubes can grow well at a relatively high temperature of 37 °C and monitor the adaptions of the pollen tubes on its lipidome, transcriptome and metabolome.

## Results

### Tobacco *in vitro* pollen tube growth is not inhibited by elevated temperatures

The *Nicotiana tabacum* genome has been sequenced, tobacco flowers are large and rich in pollen that germinates and grows tubes effectively *in vitro* (Read, Clarke and Bacic, 1993; Edwards *et al.*, 2017) making it a suitable species to use in a study on pollen tubes. First, we tested if tobacco pollen tubes also grow at higher temperatures, and thereby pose a possible model to study successful heat adaptation. For this, we analysed pollen tube elongation under five different conditions (Figure 1A) with or without HS: 3 h of growth at room temperature (22 °C, RT 3 h), 6 h of growth at RT (RT 6 h), 3 h growth at RT and then 3 h growth under HS (HS 3+3 h), 3 h growth at RT and then 6 h growth under HS (HS 3+6 h), and 3 h growth at RT followed by 3 h growth under HS and finally 3 h at RT for heat stress relief (HSR 3+3+3 h). Pollen tubes grown at RT for 3 h reached an average length of 0.46 mm and after 6 h they more than double their lengths, reaching 1.1 mm. If shifted to HS after 3 h of RT growth, pollen tube length after 6 h only reached 0.83 mm. This is a growth reduction of around 38 % in comparison to RT 6 h, indicating that while pollen tubes can still grow at this temperature their growth is hampered. If subjected to prolonged HS of 6 h, tubes averaged a length of 1.27 mm while HS relieved pollen tubes grew to 1.19 mm.

**Figure 1:**
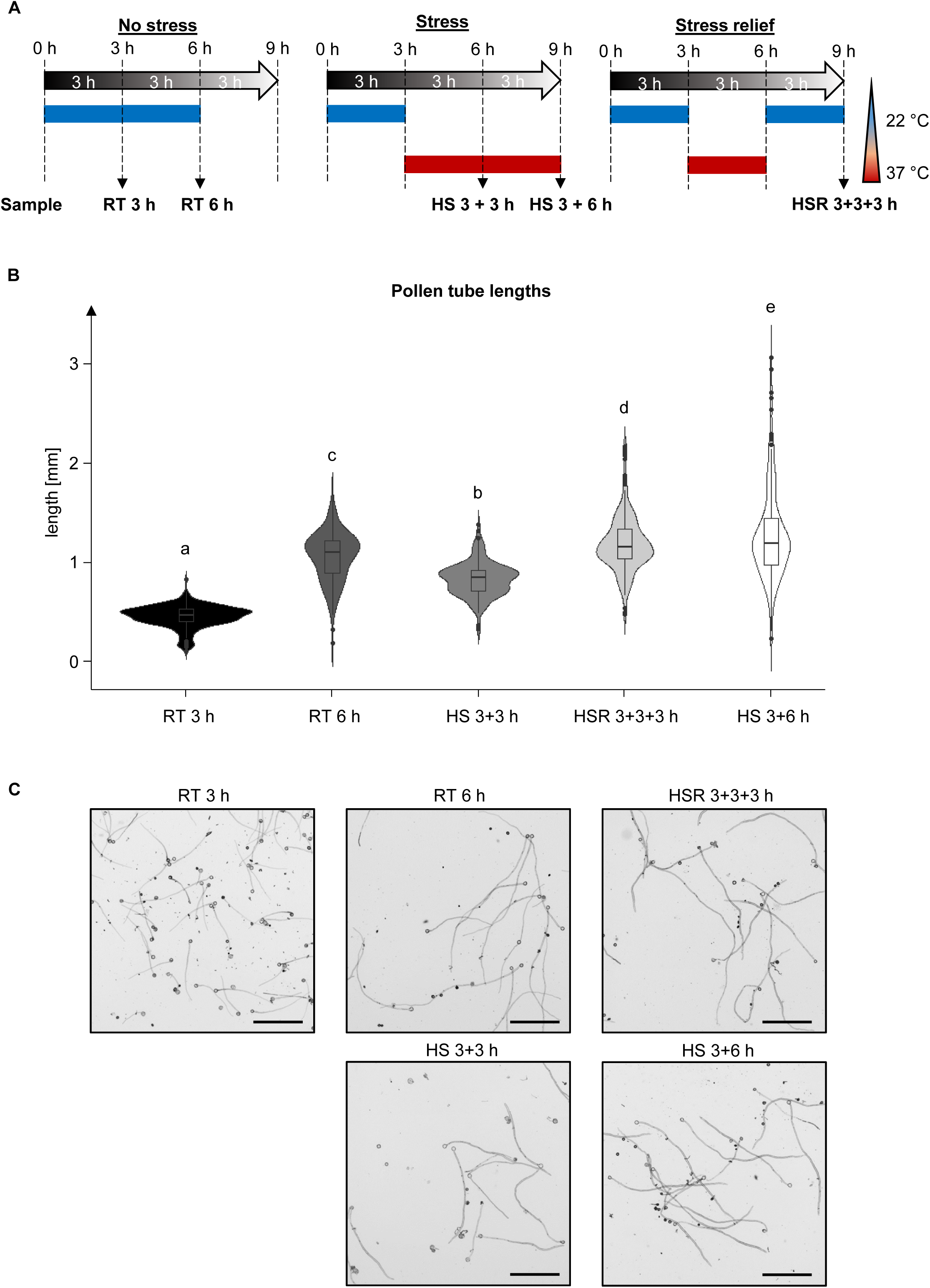
Pollen tubes were subject to different temperature regimes. **A** Schematic depiction of the experimental setup. Pollen tubes were grown under five different conditions: either unstressed for 3 h or 6 h at room temperature (22°C; RT 3 h, RT 6 h); at RT for 3 h followed by either 3 or 6 h of heat stress (37°C; HS 3+3 h, HS 3+6 h); or for 3 h at RT, followed by 3 h HS at 37 °C and then another 3 h at RT to assay heat stress relief (HSR 3+3+3 h). **B** Violin plot of measured pollen tube lengths under the different conditions. Boxplots are shown within the diagram. ANOVA was performed, followed by Post-hoc Tukey analysis. Results are presented as compact letter display of all pair-wise comparisons. n = 183 – 201. **C** Images of pollen tubes grown under the indicated conditions. Bars = 0.5 mm.

### Relative glycerophospholipid and galactoglycerolipid composition changes under heat stress

HS usually induces membrane remodelling in order to maintain membrane integrity. Sustaining pollen tube growth under elevated temperatures thus likely requires lipidomic adaptations, too. To assay such a remodelling, pollen tubes were grown under the described temperature regimes (Figure 1A). Lipids were extracted from the tubes and attached medium. Then, glycerolipids (including neutral glycerolipids and glycolipids), glycerophospholipids, sphingolipids and sterol lipids (sterol conjugates) were analysed by UPLC-nanoESI-MS/MS; free sterols were analysed by GC-MS. In total, 383 molecular species from 26 lipid subclasses were detected and relative lipid subclass-specific profiles were calculated as percent values of the individual subclasses. Abundances of these subclasses were expressed by normalization of the absolute peak area to the values after 3 h of RT growth (Supplemental Datasets S1-S6 and S9-S11). Digalactosylmonoacylglycerol, monogalactosylmonoacylglycerol, sulfoquinovosylmonacylglycerol, sulfoquinovosyldiacylglycerol, phosphatidylinositol-monophosphate (PIP), phosphatidylinositol-bisphosphate (PIP_2_) and lysophosphatidylserine could not be detected in any of the samples. Ceramide-phosphate (CerP), glucuronosylinositolphosphoceramide (GlcA-IPC), hexosyl-GlcA-IPC and hexosylhexosyl-GlcA-IPC were also not found.

HS had only mild effects on relative abundances of most membrane lipid classes (Figure 2A). For phosphatidic acid (PA), phosphatidylcholine (PC), phosphatidylinositol (PI) and phosphatidylethanolamine (PE), only subtle or no changes were observed when the tubes were grown for an additional 3 h at RT or 3 h under HS (total growth of 6 h). After a total growth of 9 h, the relative amounts of PC and PE increased, but accumulation was slightly reduced in heat-stressed tubes when compared to stress-relieved tubes. Stronger relative increases were observed for the other membrane lipids, especially so after prolonged pollen tube growth. Particularly DGDG, PS and TG showed strong relative increases upon heat treatment (Figure 2A). The effects on the synthesis and breakdown of diacylglycerol (DG), an intermediate of several lipid pathways, were less pronounced (Figure 2A).

**Figure 2:**
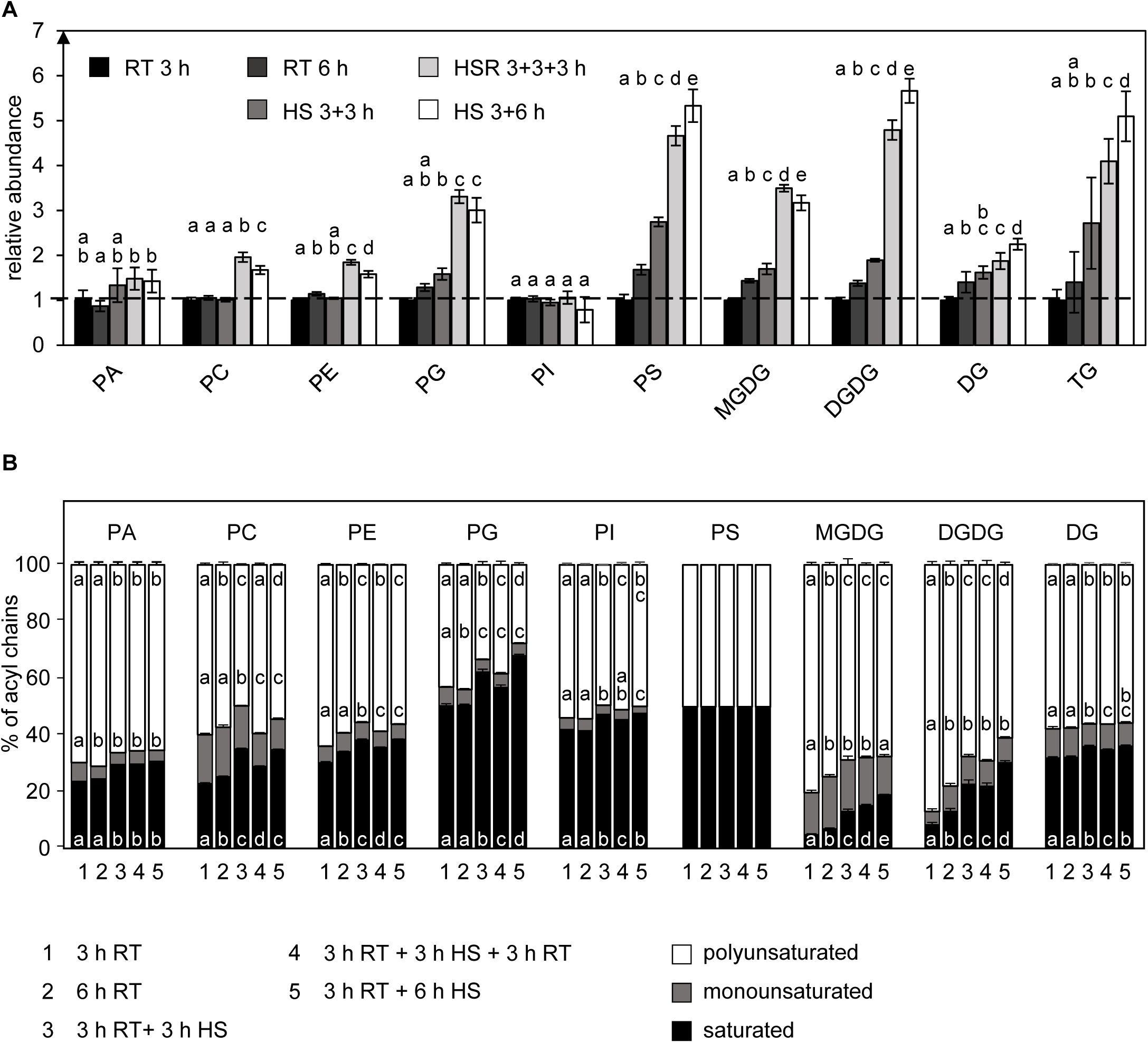
Heat stress changes lipid subclass composition and decreases saturation of acyl chains. **A** Abundances of the depicted lipid subclasses of pollen tubes grown under different temperature regimes (RT, room temperature; HS, heat stress; HSR, heat stress relief) were measured by UPLC-nanoESI-MS/MS and normalized to their respective values after 3 h RT. **B** Relative saturation of the respective lipid classes under the same temperature regimes. The levels of the molecular lipid species containing zero (saturated), one (monounsaturated) or more than one (polyunsaturated) double bonds in their fatty acid residues were summed per lipid subclass and converted into relative molar proportions. n=5. Error bars represent standard deviation. For statistical analysis, ANOVA was performed, followed by Post-hoc Tukey analysis. Results are presented as compact letter display of all pair-wise comparisons. PA, phosphatidic acid; PC, phosphatidylcholine; PE, phosphatidylethanolamine; PG, phosphatidylglycerol; PI, phosphatidylinositol; PS, phosphatidylserine; MGDG, monogalactosyldiacylglycerol; DGDG, digalactosyldiacylglycerol; DG, diacylglycerol; TG, triacylglycerol.

Looking at overall saturation levels, the relative proportion of saturated acyl chains increased in all membrane glycerolipid subclasses except for PS and DG, while the relative abundance of polyunsaturated acyl chains decreased (Figure 2B). Shifting the tubes back to RT partly led to a reversion of the effects in PC, PE, and PG while relative levels in MGDG and DGDG remained approximately constant.

Regarding the acyl chain composition (Figure 3), a relative increase in most 16:0- and 18:0-containing glycerophospholipid species was observed after 3 h of HS, while species without saturated acyl chains generally decreased in relative abundance. If pollen tubes were shifted back to RT, most species containing 16:0 and 18:0 decreased again; while lipids containing no saturated acyl chains increased.

**Figure 3:**
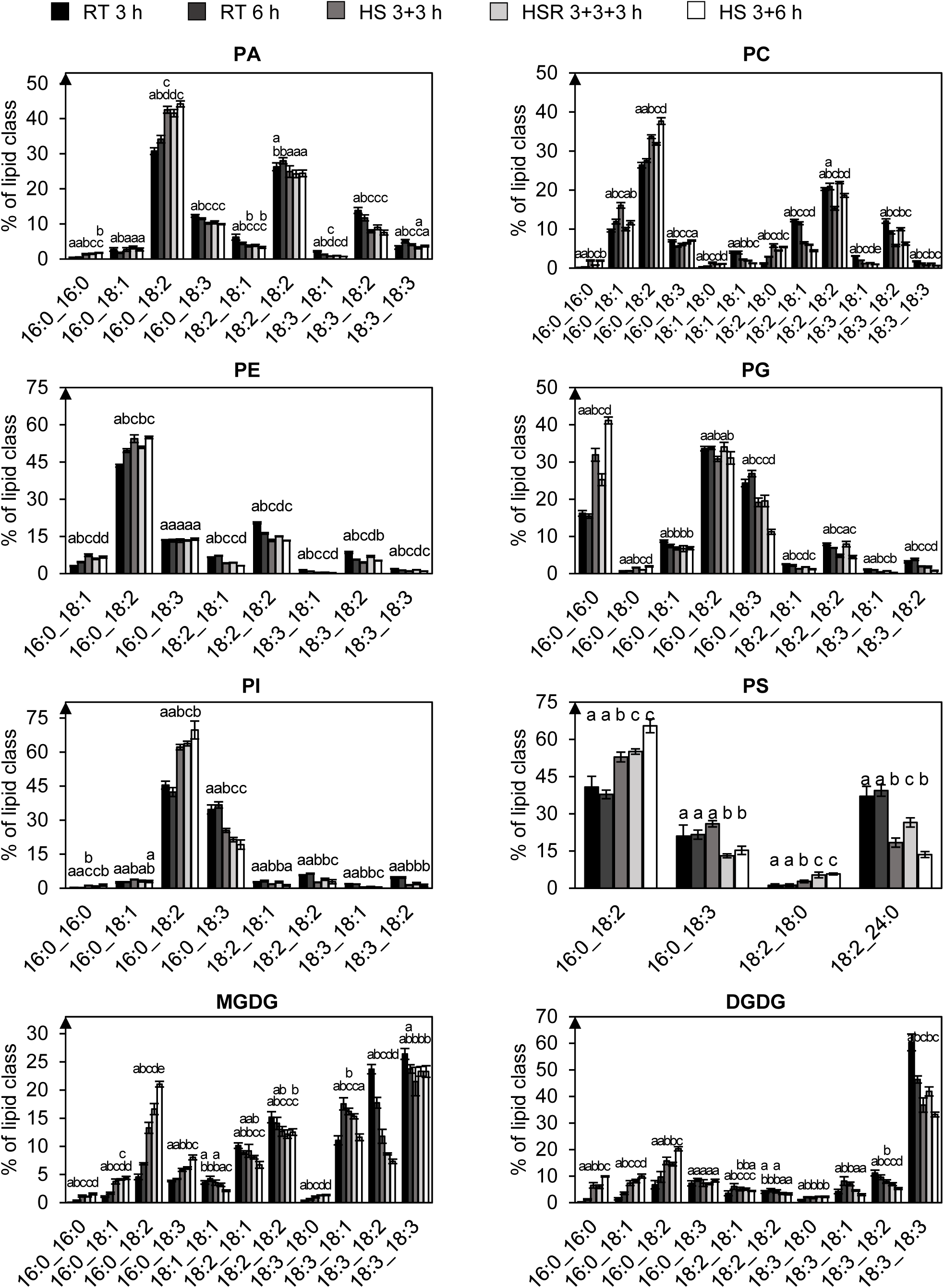
Glycerophospholipid and galactolipid species with saturated acyl chains increase under heat stress. The depicted lipid subclasses from pollen tubes grown under different temperature regimes (RT, room temperature; HS, heat stress; HSR, heat stress relief) were investigated. Lipid subclass profiles were obtained by UPLC-nanoESI-MS/MS. Values are given as mol % of all detected lipid species per subclass, only major lipid species (> 1% of total) are depicted. n=5. Error bars represent standard deviation. For statistical analysis, ANOVA was performed, followed by Post-hoc Tukey analysis. Results are presented as compact letter display of all pair-wise comparisons. PA, phosphatidic acid; PC, phosphatidylcholine; PE, phosphatidylethanolamine; PG, phosphatidylglycerol; PI, phosphatidylinositol; PS, phosphatidylserine; MGDG, monogalactosyldiacylglycerol; DGDG, digalactosyldiacylglycerol.

Monoacylglycerophospholipids that might negatively affect membrane rigidity (Henriksen *et al*., 2010) were strongly reduced under HS (Figure 4A). The relative proportion of most monoacylglycerophospholipids containing 16:0 also increased after 3 h of HS (Figure 4B-F). The relative amounts of 18:2- and 18:3-containing species on the other hand declined leading in conclusion to a higher saturation (Figure 4G). These effects were however not enhanced under continuous growth at 37 °C.

**Figure 4:**
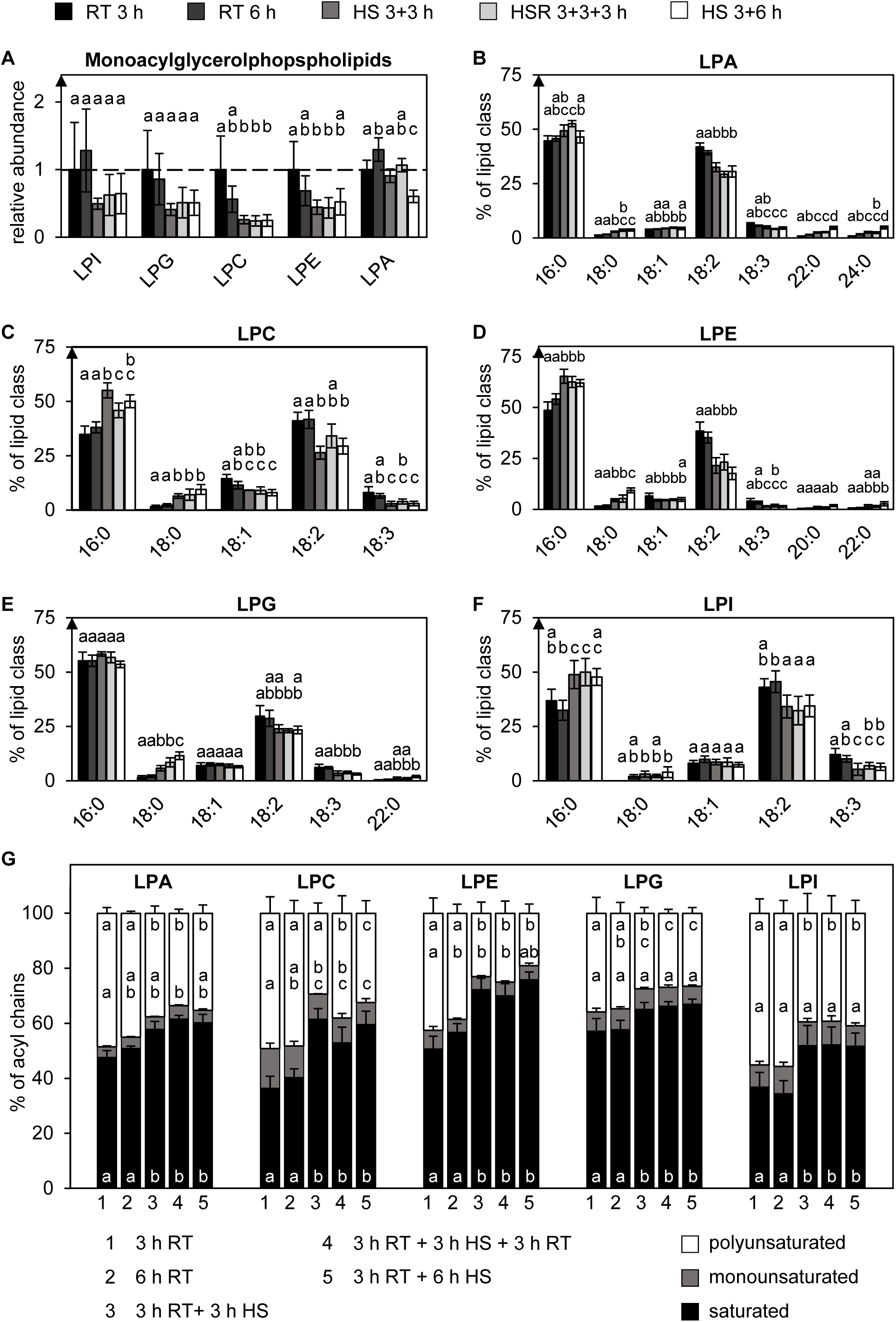
Monoacylglycerphospholipid species with saturated acyl chains increase under heat stress. The depicted lipid subclasses from pollen tubes grown under different temperature regimes (RT, room temperature; HS, heat stress; HSR, heat stress relief) were investigated. **A** Relative abundances of the lipid subclasses were measured by UPLC-nanoESI-MS/MS and normalized to their respective values after 3 h RT. **B-F** Molecular species within the individual subclasses are given as mol % of all detected lipid species per subclass. Only major lipid species (> 1% of total) are depicted. **G** Relative saturation of the respective lipid classes under the same temperature regimes. The abundance of the molecular lipid species containing zero (saturated), one (monounsaturated) or more than one (polyunsaturated) double bonds in their acyl residues were summed per lipid subclass and converted into relative proportions. n=5. Error bars represent standard deviation. For statistical analysis, ANOVA was performed, followed by Post-hoc Tukey analysis. Results are presented as compact letter display of all pair-wise comparisons. LPA, lysophosphatidic acid; LPC, lysophosphatidylcholine; LPE, lysophosphatidylethanolamine; LPG, lysophosphatidylglycerol; LPI, lysophosphatidylinositol.

The general trend observed in MGDG and DGDG is the same as for glycerophospholipids: 16:0-containing lipid species increased, while 18:3-containing lipid species decreased. Stress relief partially reversed the observed effects again for some species but not for all (Figure 3).

Fewer changes were observed in the acyl chain composition of the glycerolipids DG and TG (Figure 5A). DG profiles remained more or less constant under all tested conditions. For TG, some changes over time but almost no changes in response to HS were observed.

**Figure 5:**
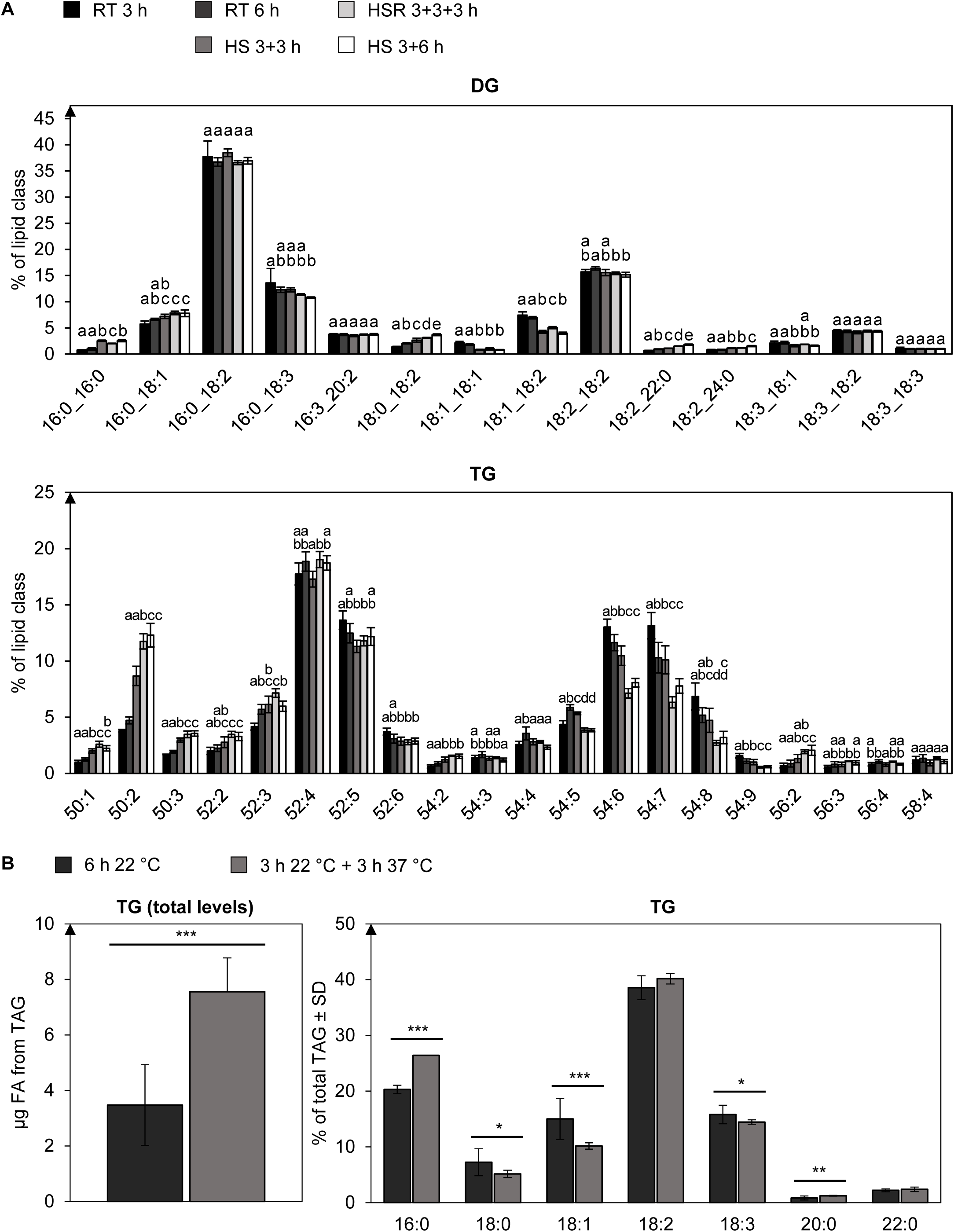
Triaclglycerol accumulates under heat stress and becomes more saturated. **A** The depicted lipid subclasses from pollen tubes grown under different temperature regimes (RT, room temperature; HS, heat stress; HSR, heat stress relief) were investigated. Lipid subclass profiles were measured by UPLC-nanoESI-MS/MS. Values are given as mol % of all detected lipid species per subclass. Only major lipid species (> 1% of total) are depicted. For statistical analysis, ANOVA was performed, followed by Post-hoc Tukey analysis. n=5. Error bars represent standard deviation. Results are presented as compact letter display of all pair-wise comparisons. DG, diacylglycerol; TG, triacylglycerol. **B** Absolute quantification of TG from heat stressed and non-stressed pollen tubes per 1 mg of dry pollen. Also shown is the relative contribution of the respective fatty acids to the total TG amount. Measurements were performed with GC-FID and quantified as peak areas. n=5. Error bars represent standard deviation. *** p < 0.005; ** p < 0.001; * p < 0.05; determined by Student’s *t*-test.

To further validate the relative increase of TG levels under HS (Figure 2A), and to get information on absolute TG levels, TG content were also quantified by GC-FID (Figure 5B). Heat-stressed pollen tubes (3 h RT + 3 h HS) produce more than twice the absolute amount of TG as control pollen tubes (6 h RT) did.

### Pollen tube sphingolipid abundance increases under heat stress

Sphingolipids are a highly diverse class of lipids. All sphingolipids consist of a sphingoid base (SPB) that can be subject to different modifications (Michaelson *et al.*, 2016). If the SPB is linked to a 14-30 carbon acyl chain via an amide bond, a ceramide is formed. Ceramides can be linked to a head group, consisting either of a hexose (giving rise to the subclass of hexosylceramides; HexCers) or a phosphate attached to an inositol. This inositol can then be further linked to glucuronic acid forming the core structure of glycosylinositolphosphoceramides (GIPCs), which can carry additional sugar residues. Both, the SPBs and the fatty acids can be further modified by double bonds or hydroxyl groups, SPBs and ceramides can also be phosphorylated to either SPB-phosphates (SPBPs) or Cer-Ps (Figure 6A).

**Figure 6:**
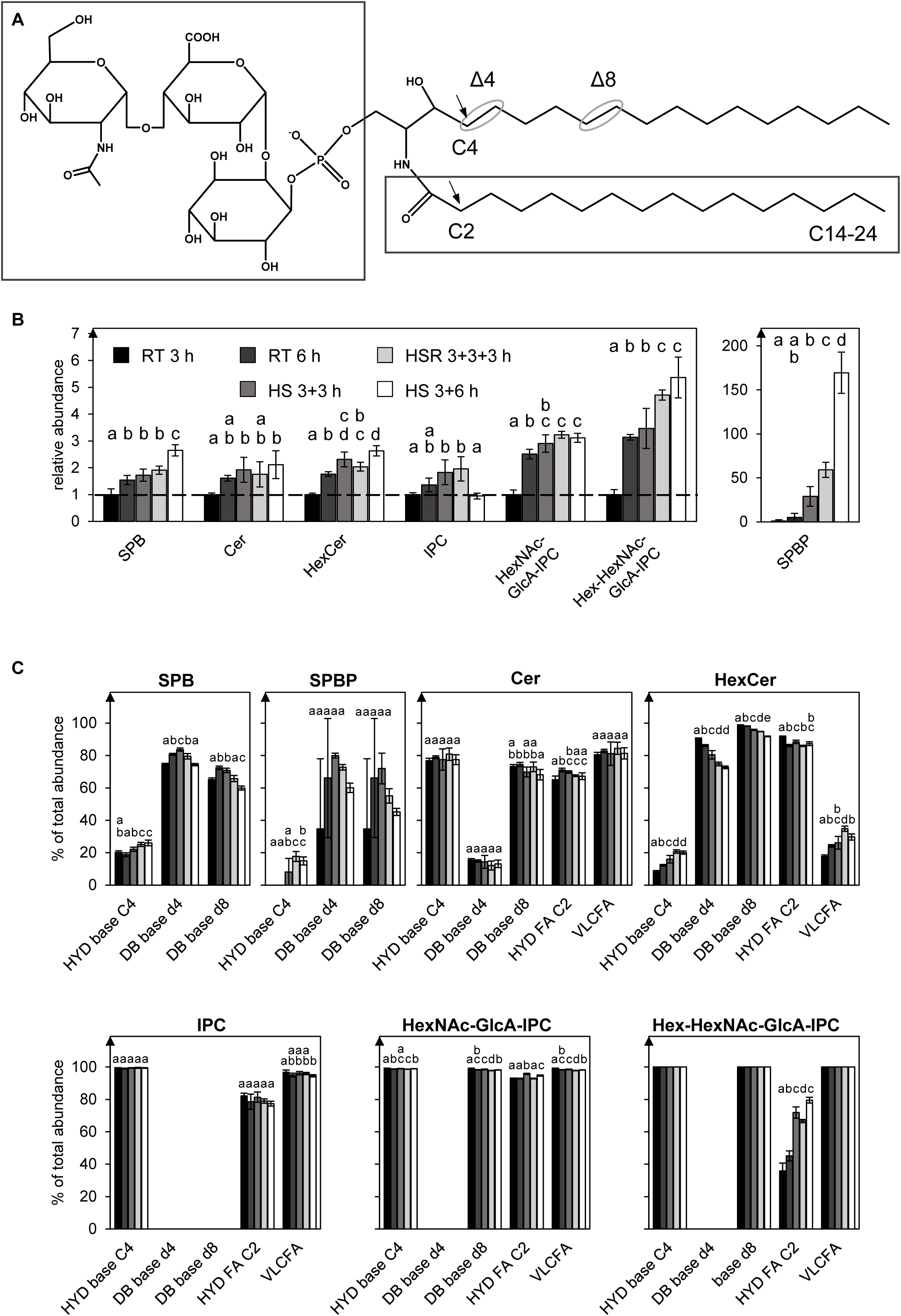
Sphingolipid composition and modifications are affected by heat stress. **A** Sphingolipids consist of a sphingoid base (SPB) bound to an acyl chain of variable length (right box) forming a ceramide. They can carry a head group (left box) consisting of a hexose (hexosylceramide, HexCer), a phosphate (for example SPBP), a phosphate fused to inositol (inositolphosphoceramide, IPC) that can be further linked via glucuronic acid (GlcA) to additional sugars as depicted for a hexosyl-*N*-acetylhexosaminyl-GlcA-IPC (HexNAc-GlcA-IPC) that can also be linked to a further hexose (Hex-HexNAc-GlcA-IPC). In addition, sphingolipids can be modified at the SPB by a double bond in the d4 and d8 position or (mutually exclusive with the d4 double bond) a hydroxylation at the C4 carbon atom. Another variable modification that can be detected is a hydroxylation at the C2 of the acyl chain. **B** The depicted sphingolipid subclasses from pollen tubes grown under different temperature regimes (RT, room temperature; HS, heat stress; HSR, heat stress relief) were investigated. Abundances of the lipid subclasses were measured by UPLC-nanoESI-MS/MS and normalized to their respective values after 3 h RT. **C** For each sphingolipid subclass, the percentage-wise occurrence of modifications was determined: HYD base C4, hydroxylation at C4 of the SPB; DB base d4, double bond at the Δ4 position of the SPB; DB base d8, double bond at the Δ8 position of the SPB; HYD FA C2, hydroxylation at C4 of the acyl chain; VLCFA, acyl chain derived from a very long chain fatty acid (C ≥ 20). n=5. Error bars represent standard deviation. For statistical analysis, ANOVA was performed, followed by Post-hoc Tukey analysis. Results are presented as compact letter display of all pair-wise comparisons.

Overall, 96 sphingolipid species from 7 lipid subclasses were identified in the samples (Supplemental Datasets S4-S6). The abundance of sphingolipids in tobacco pollen tubes increased over time (Figure 6B), and HS led to an additional increase of all sphingolipid subclasses. Especially the levels of SPBP (5.8-fold increase when comparing HS 3+3 h to RT 6 h, and 2.8-fold when comparing extended HS to HSR) and HexCer (30 % and 31 % increase, respectively) were elevated.

Sphingolipids of Arabidopsis pollen were previously described to be distinct from sporophytic tissues in having HexCer species that predominantly carry a double bond at the C4 position of the SPB instead of a hydroxy group (Luttgeharm *et al*., 2015). Also in this study, after 3 h of growth, this double bond was detected in at least 90 % of the HexCer species, in 74 % of the SPB species and in 15 % of the ceramide species (Figure 6C; for some lipid species the position of the hydroxyl group could not unambiguously be determined). Interestingly, the occurrence of this modification was slightly reduced under HS. Instead, the relative abundance of species containing a hydroxyl group at this position increased (by 31 % when compared to HS 3+3 h to RT 6 h). Another major change observable under HS was the increased hydroxylation at the C2 of the acyl chain of hexosyl-*N*-acetylhexosaminyl-GlcA-IPCs (Hex-HexNAc-GlcA-IPCs), which was detected in 45 % of all Hex-HexNAc-GlcA-IPC species after 6 h growth at RT but increased to 72 % and 80 % under HS and extended HS, respectively.

### Levels and composition of free sterols and sterol conjugates are altered under heat stress conditions

Similar to sphingolipids, sterol lipids are important structural lipid constituents of membranes, especially the plasma membrane, and are important for membrane microdomain (lipid raft) formation. Through this, they are involved in diverse cellular processes, including polar cell growth (Simons and Ikonen, 1997; Beck *et al.*, 2007; Simon-Plas *et al.*, 2011; Mamode Cassim *et al.*, 2019).

The profile of free sterols in pollen from various species differs greatly from that in leaves or other sporophytic tissues, and it was shown that the sterol synthesis pathway in growing tobacco pollen tubes is truncated with only two species being *de novo* synthesised: presumably methylenepollinastanol and its precursor cycloeucalenol (Villette *et al.*, 2015; Rotsch *et al.*, 2017). Relative abundance of these sterol species increases during pollen tube growth (Rotsch *et al.*, 2017), as can be seen in the presented data (Figure 7, Supplemental Datasets S7-S8). Especially cycloeucalenol accumulated strongly under HS and less at RT. After HSR, increased accumulation ceased and levels remained constant.

**Figure 7:**
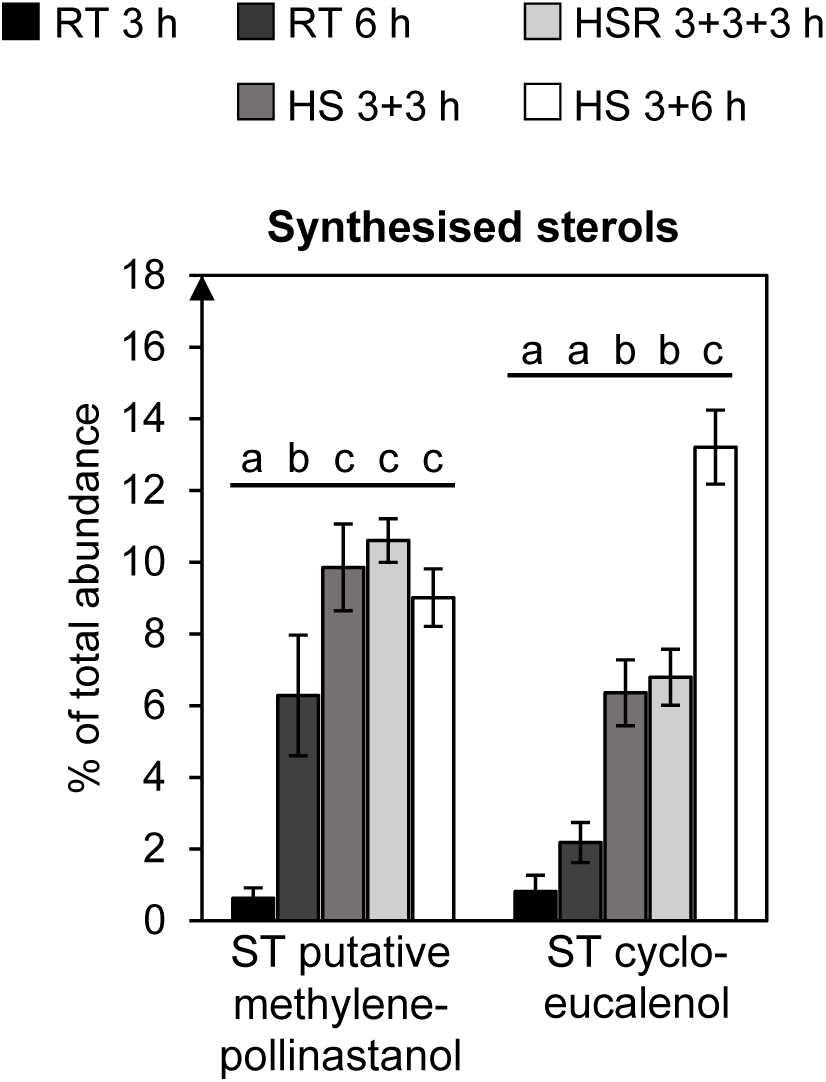
The relative amounts of newly synthesised free sterol species increase under heat stress. Only two sterol species are newly synthesised in growing tobacco pollen tubes, a sterol with an exact mass of 412,37 Da that is putatively methylenepollinastanol, and cycloeucalenol. Presented are relative abundances of the respective sterol based on the total ion current as measured by GC-MS, pollen tubes were grown under different temperature regimes (RT, room temperature; HS, heat stress; HSR, heat stress relief). For statistical analysis, ANOVA was performed, followed by Post-hoc Tukey analysis. Results are presented as compact letter display of all pair-wise comparisons in increasing order.

Also the relative levels of sterylglycoside (SG) and acylsterylglycoside (ASG) were strongly increased by 84 % and 42 %, respectively when comparing 3 h of HS to the control treatment. Sterol ester (SE) species on the other hand decreased in their relative levels (Figure 8, Supplemental Datasets S10-S11). Prolonged heat treatment did not lead to a further strong increase or decrease of sterol conjugates. However, HSR caused SE levels to increase again.

**Figure 8:**
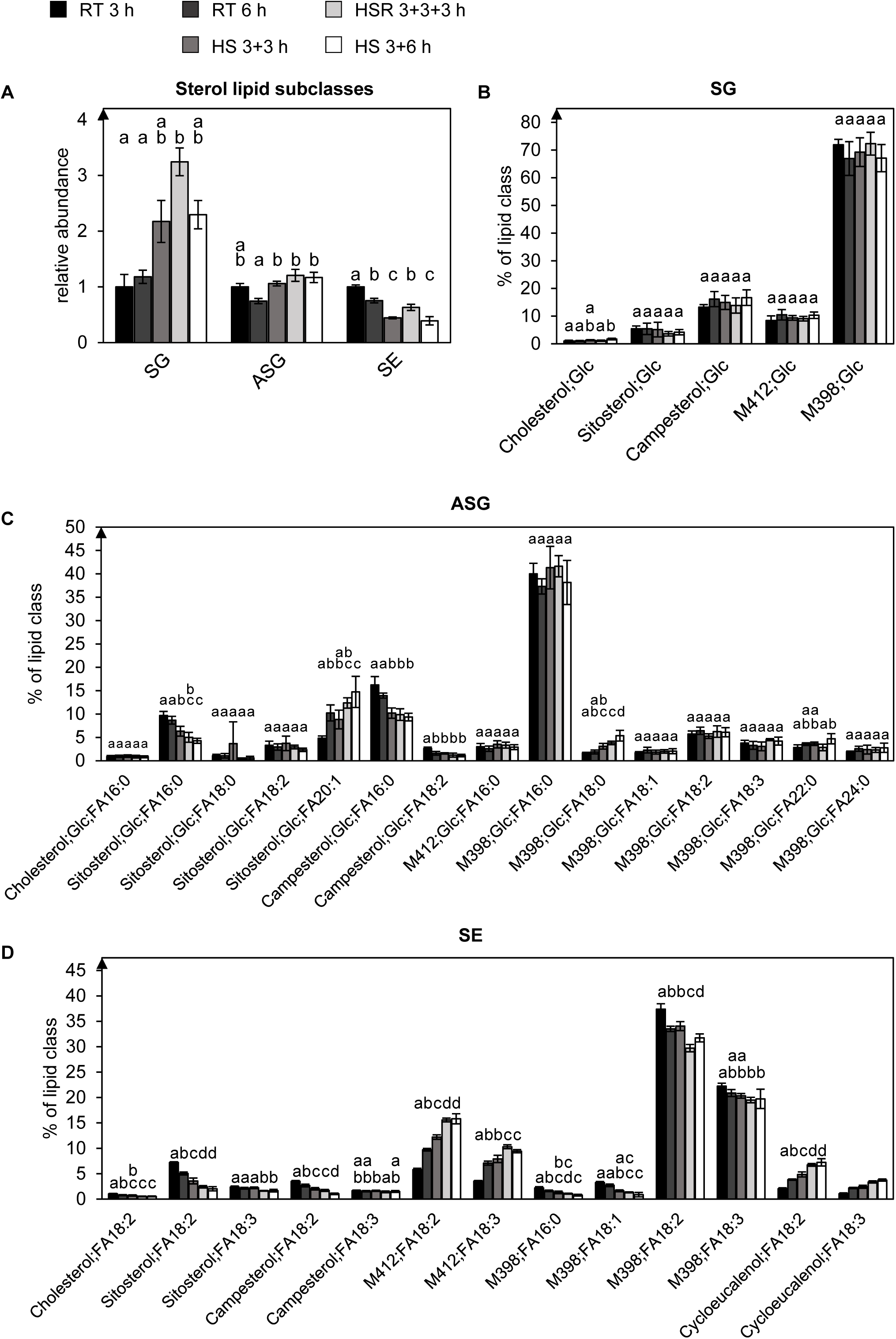
Sterylglycosides accumulate under heat stress, while sterol esters decrease. **A** The depicted sterol lipid subclasses from pollen tubes grown under different temperature regimes (RT, room temperature; HS, heat stress; HSR, heat stress relief) were investigated. Abundances of the lipid subclasses were measured by UPLC-nanoESI-MS/MS and normalized to their respective values after 3 h RT. **B-D** Molecular species within the individual subclasses are given as mol % of all detected lipid species per subclass. Only major lipid species (> 1% of total) are depicted. M398 could represent methylenecholesterol or Δ5,24-ergostadienoland, and M412 stigmasterol, the putative methylenepollinastanol, isofucosterol or 24-methylenelophenol. For statistical analysis, ANOVA was performed, followed by Post-hoc Tukey analysis. Results are presented as compact letter display of all pair-wise comparisons in increasing order.

### Several central metabolites increase under HS

In our study, we also extracted hydrophilic metabolites from pollen tubes grown under the five temperature regimes (Figure 9A). As pollen tubes cannot be easily washed, some liquid attached to the tubes that could contain secreted metabolites was included in the measurements. We identified 51 metabolites by GC-MS that are mostly part of central metabolism including organic acids, amino acids and sugars (Supplemental Datasets S12-S13). In addition, 17 so far unidentified markers were found. Values of all time points were normalised to the values after 3 h of RT. While most metabolites accumulated during prolonged pollen tube growth, not all of these were affected by HS. Sugar levels were only mildly altered with the exception of the 7-carbon sugar sedoheptulose that increased 3-fold, and sucrose that increased 2- fold (Figure 9B). Among the organic acids, an increase in 2-isopropylmalate, an intermediate in leucine biosynthesis, and 2-oxoglutarate was observed (Figure 9C). Interestingly, fumarate and malate were not affected by HS but strongly increased after stress relief. Most amino acids increased during HS, including the non-codogenic amino acids β-lactate, γ-amino butyrate (GABA) and pipecolate (Figure 9D).

**Figure 9:**
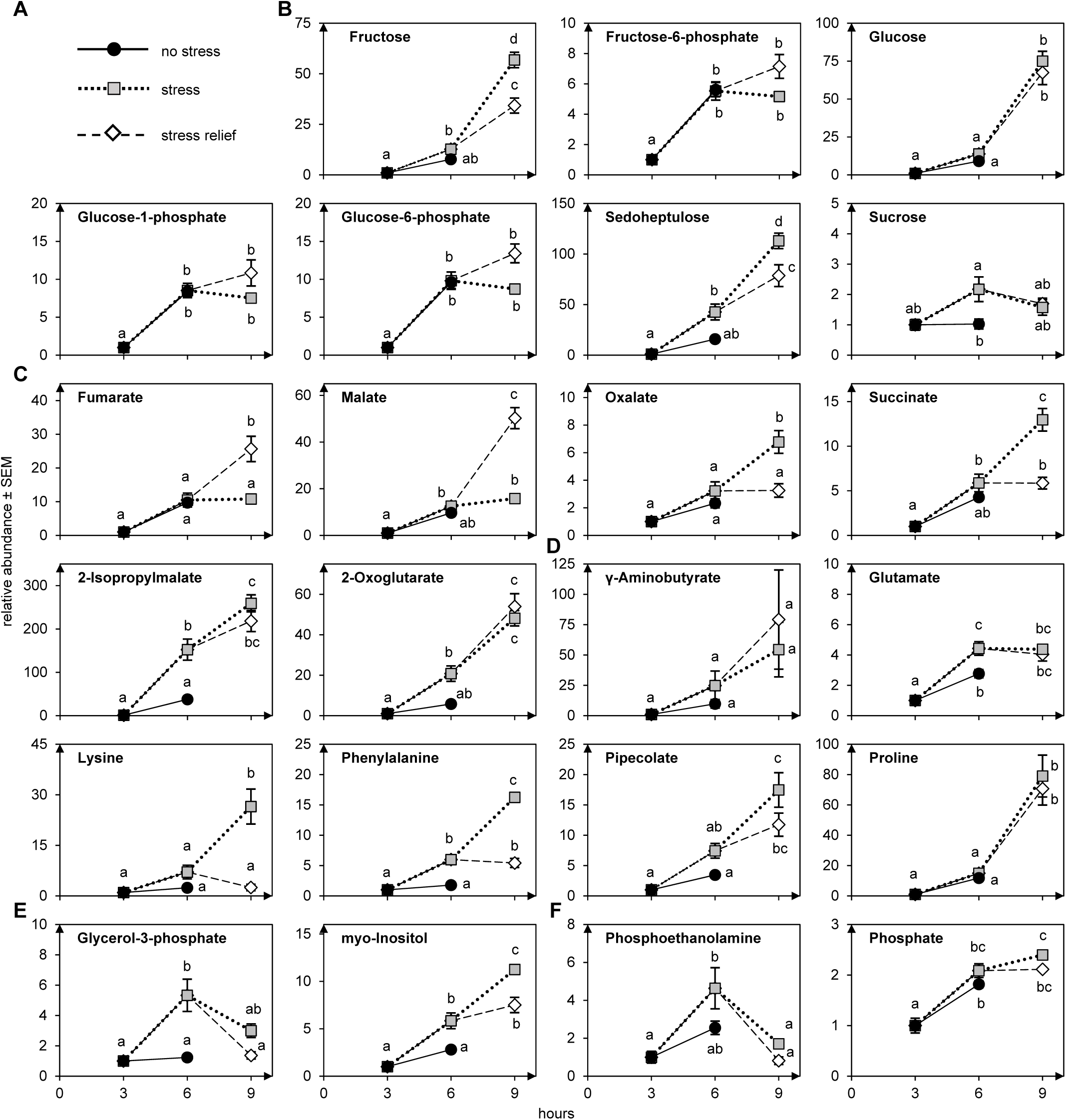
The abundance of several metabolites is affected by heat stress. **A** Legend: black line and filled black circles: no stress (RT 3h and RT 6); dotted line and grey squares: heat stress (HS 3+3h and HS 3+6h); dashed lines and white rhombus: heat stress relief (HSR 3+3+3h). Shown are a selection of detected sugars (**B**), a selection of detected organic acids (**C**), a selection of detected amino acids (**D**), a selection of detected polyols (**E**) and others (**F**). n=5. Metabolites are normalized to the value at 3 h. Error bars represent standard deviation. For statistical analysis, ANOVA was performed, followed by Post-hoc Tukey analysis. Results are presented as compact letter display of all pair-wise comparisons in increasing order.

### Heat stress leads to strong alteration on the transcriptome level

To assess whether the different adaptations are reflected on a transcriptional level, transcriptome analyses of pollen tubes grown for 3 or 6 h at RT or 3 h at RT followed by 3 h at HS were performed by RNA sequencing (Supplemental Dataset S14). The corresponding PCA plot is shown in Figure 10A.

**Figure 10:**
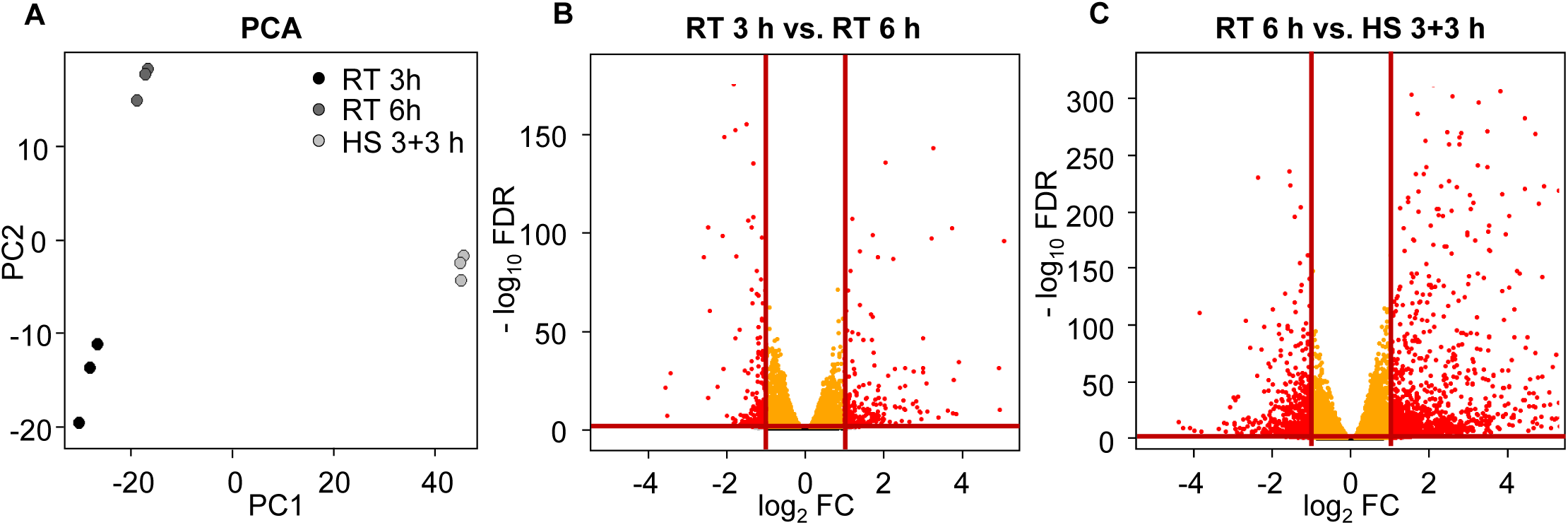
Analysis of transcriptome data. mRNA levels from pollen tubes grown under different temperature regimes (RT, room temperature; HS, heat stress) were investigated by RNA sequencing. **A** Principle component analysis (PCA) of RT 3 h, RT 6 h and HS 3+3 h. **B and C** Volcano plots of RT 3h vs. RT 6h and RT 6h vs. HS 3+3 h, respectively. Red lines indicate threshold for differential expression (abs. log_2_ FC > 1, red dots and FDR < 0.005, orange dots).

Overall, 26743 genes were detected, 24,013 of which with more than 0.5 counts per million (CPM) in at least 2 libraries and 21,241 of these could be assigned a UniProt protein ID (Table 1). The respective protein sequences were then blasted against the TAIR 10 Arabidopsis protein library, and 19,798 tobacco proteins had Arabidopsis homologues with an Expect value (E-value) < 10^-5^. The respective Arabidopsis protein identifiers were later used for functional annotation.

**Table 1:** Number of detected and annotated genes in the respective analysis.

| Analysis | Detected genes | UniProt ID | Arabidopsis homolog |
| --- | --- | --- | --- |
| RT 3 h vs. RT 6 h | 23,043 | 20,459 | 19,162 |
| RT 6 h vs. HS 3+3 h | 23,053 | 20,445 | 19,779 |
| total | 24,013 | 21,241 | 19,798 |
| **total unfiltered** | 26,743 | - | - |

Number of detected genes in total and in the two comparisons and number of total detected genes (unfiltered). For DE analyses, genes were filtered for a CPM < 0.5 in at least two of the respective analysed libraries. Genes were annotated a UniProt identifier (UniProt ID), some genes could not be annotated, number of annotated genes is given. Annotated Tobacco proteins were blasted against Arabidopsis to find closest homologs, only hits with E- value < 10<sup>-5</sup> were considered. Number of genes with an Arabidopsis protein homolog hit is given.

In the analysis of pollen tubes grown for three versus six hours at RT, fold-changes (FCs) were calculated. 561 genes were differentially expressed (defined as log_2_FC > 1; FDR ≤ 0.005), 306 of which were upregulated and 255 of which were downregulated (Figure 10B, Supplemental Dataset S15).

In the comparison of heat-stressed versus non heat-stressed pollen tubes, 2,383 genes were differentially expressed, 1,570 of which were upregulated and 813 of which were downregulated (Figure 10C, Supplemental Dataset S16). For analyses of gene functions, we further explored the second comparison making use of the functional annotation of the Arabidopsis genome and proteome.

### GO-term analysis gives first insights

For gene ontology (GO) analysis, the GO terms assigned to the annotated Arabidopsis homologues were analysed. For the analysis of RT 6 h versus HS 3+3 h, 20,370 genes were assigned with 5994 different GO terms. Reads per kilo base per million mapped reads (RPKM) values of all genes belonging to the same GO term were summed up and averaged for the respective condition to calculate log_2_FCs.

A selection of the most changed GO terms between RT 6 h and HS 3+3 h are presented in Table 2, grouped by sub-ontology (for complete lists see Supplemental Dataset S17-20). Among them are several heat shock-related GO terms containing many upregulated genes and only few downregulated genes. The terms include “functions in Hsp90 protein binding”, “has unfolded protein binding”, and “involved in response to heat”. Another strongly changed GO term is “involved in response to reactive oxygen species”. Other changed GO terms like “has pre-mRNA 3’-splice site binding” and “involved in positive regulation of mRNA splicing, via spliceosome” might hint at a role of alternative splicing under HS, which has been shown before, e.g. in tomato pollen (Keller *et al.*, 2017). Interestingly, several changed GO terms suggest that auxin metabolism and/or signalling might be affected (“has auxin receptor activity”, “involved in auxin catabolic process”, and “involved in regulation of auxin biosynthetic process”).

**Table 2:**
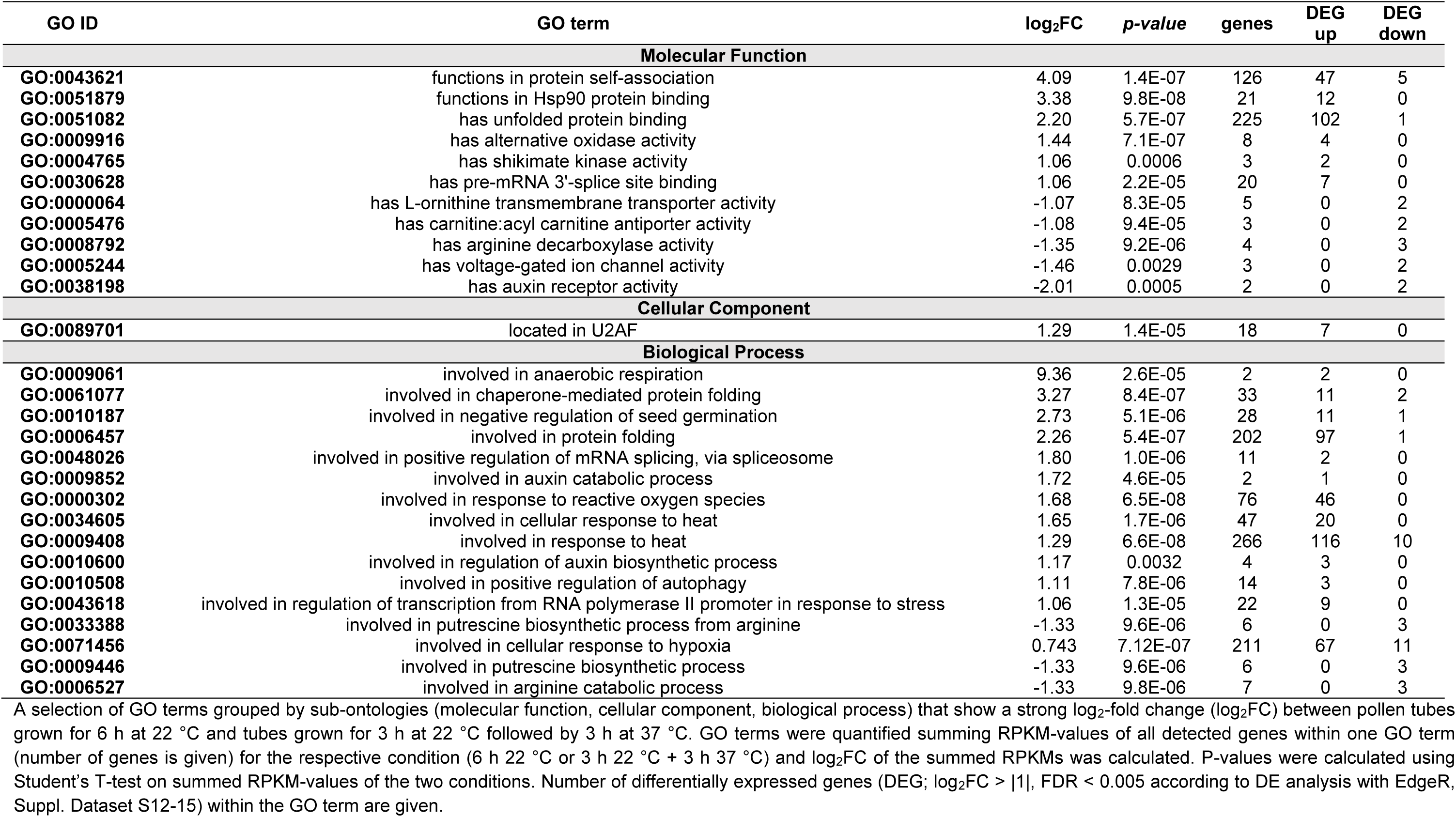
List of selected GO-terms, their IDs, log_2_-fold changes (6 h 22 °C vs. 3 h 22 °C + 3 h 37 °C), p-values, the number of detected genes within the GO-term and the number of differentially expressed genes (up- and downregulated) within the term.

### Transcripts of transcriptional regulators are strongly affected

The GO term analysis revealed the term “has transcription coactivator activity” to be 3.2-fold upregulated. The terms “functions in transcription regulatory region DNA binding” and “has DNA-binding transcription factor activity” showed 34 of 395 and 72 of 654 differentially upregulated, but also 31 and 15 downregulated genes, respectively. The total transcript abundance within these terms was, however, only little changed (Supplemental Dataset S19). To take a closer look at putative transcriptional regulators important for HS adaptation, transcript data were mined for genes with homology to Arabidopsis transcriptional regulators according to the Arabidopsis Plant Transcription Factor Database with 2192 gene entries (Pérez-Rodríguez *et al.*, 2010, Supplemental Dataset S21). Overall, 1319 tobacco genes could be assigned to transcriptional regulators (117 up- and 42 differentially downregulated; Table 3, Supplemental Dataset S22). The highest proportion of upregulated genes was found among the heat shock transcription factors (HSF, 9 out of 19) and DNA-binding protein phosphatases (DBP, 6 out of 10). Furthermore, the transcription factor families MYB-related and AP2-EREBP showed a comparably high number of upregulated genes (14 and 9, respectively).

**Table 3:** List of all detected transcription factor families with differentially expressed genes.

| Transcription factor family | Type | Arabidopsis genes | Detected genes | DEG up | DEG down |
| --- | --- | --- | --- | --- | --- |
| ABI3VP1 | TF | 57 | 27 | 1 | 1 |
| Alfin-like | TF | 7 | 10 | 0 | 1 |
| AP2-EREBP | TF | 146 | 34 | 9 | 1 |
| ARF | TF | 23 | 15 | 2 | 1 |
| ARID | Other | 10 | 17 | 1 | 1 |
| AUX/IAA | Other | 28 | 1 | 1 | 0 |
| BES1 | TF | 8 | 10 | 0 | 1 |
| bHLH | TF | 136 | 44 | 5 | 3 |
| bZIP | TF | 70 | 49 | 1 | 5 |
| C2C2-CO-like | TF | 17 | 4 | 0 | 1 |
| C2C2-GATA | TF | 29 | 13 | 2 | 0 |
| C2H2 | TF | 99 | 43 | 3 | 3 |
| C3H | TF | 68 | 90 | 7 | 1 |
| CCAAT | TF | 43 | 39 | 3 | 0 |
| Coactivator p15 | Other | 3 | 2 | 2 | 0 |
| CSD | TF | 4 | 3 | 2 | 0 |
| DBP | TF | 4 | 10 | 6 | 0 |
| EIL | TF | 6 | 8 | 2 | 0 |
| FAR1 | TF | 17 | 26 | 2 | 1 |
| FHA | TF | 17 | 20 | 3 | 0 |
| G2-like | TF | 40 | 20 | 2 | 0 |
| GNAT | Other | 32 | 34 | 1 | 0 |
| GRAS | TF | 33 | 17 | 3 | 3 |
| HB | TF | 91 | 19 | 1 | 0 |
| HSF | TF | 23 | 19 | 9 | 0 |
| LOB | TF | 43 | 14 | 1 | 0 |
| LUG | Other | 2 | 6 | 1 | 0 |
| MADS | TF | 105 | 30 | 2 | 2 |
| MBF1 | Other | 3 | 7 | 2 | 0 |
| mTERF | TF | 34 | 11 | 1 | 1 |
| MYB | TF | 147 | 55 | 2 | 2 |
| MYB-related | TF | 65 | 71 | 14 | 1 |
| NAC | TF | 104 | 26 | 3 | 0 |
| Orphans | TF | 83 | 35 | 4 | 1 |
| PHD | Other | 45 | 92 | 3 | 1 |
| Pseudo ARR-B | Other | 5 | 5 | 1 | 1 |
| RWP-RK | TF | 14 | 10 | 2 | 1 |
| SET | Other | 33 | 46 | 3 | 0 |
| Sigma70-like | TF | 6 | 6 | 3 | 0 |
| SNF2 | Other | 38 | 57 | 0 | 3 |
| SWI/SNF-BAF60b | Other | 16 | 10 | 1 | 0 |
| TCP | TF | 24 | 5 | 0 | 1 |
| Tify | TF | 15 | 9 | 0 | 1 |
| TRAF | Other | 25 | 16 | 0 | 2 |
| TUB | TF | 10 | 15 | 2 | 0 |
| ULT | TF | 2 | 4 | 0 | 1 |
| VOZ | TF | 2 | 4 | 1 | 0 |
| WRKY | TF | 72 | 22 | 3 | 1 |

List of all detected transcription factor families that contain differentially expressed genes. As reference, the Arabidopsis Plant Transcription Factor Database PlnTFDB with 2192 gene entries was used (Pérez-Rodríguez *et al.* 2010; accessed on 4<sup>th</sup> June 2020). Given are the transcription factor family names, their type (TF = transcription factor, Other = other transcriptional regulator), the number of Arabidopsis genes belonging to the respective family, the number of detected Tobacco genes putatively belonging to the respective family and the number of differentially expressed genes (DEG; $\log_2FC > |1|$ , $FDR < 0.005$ ) within the family.

### Transcripts related to lipid metabolism are barely affected by heat stress

To find out whether lipid adaptations are mirrored on a transcriptional level, a list of 746 Arabidopsis genes with a role or putative role in glycerolipid, sphingolipid and sterol metabolism was compiled from several sources (see method section and Supplemental Dataset S23). Comparing our transcript data to this list, 713 tobacco transcripts were assigned a putative function in lipid metabolism (Supplemental Dataset S24). Of these, only 43 genes were differentially expressed, 20 of which were up- and 23 downregulated (Table 4).

**Table 4:**
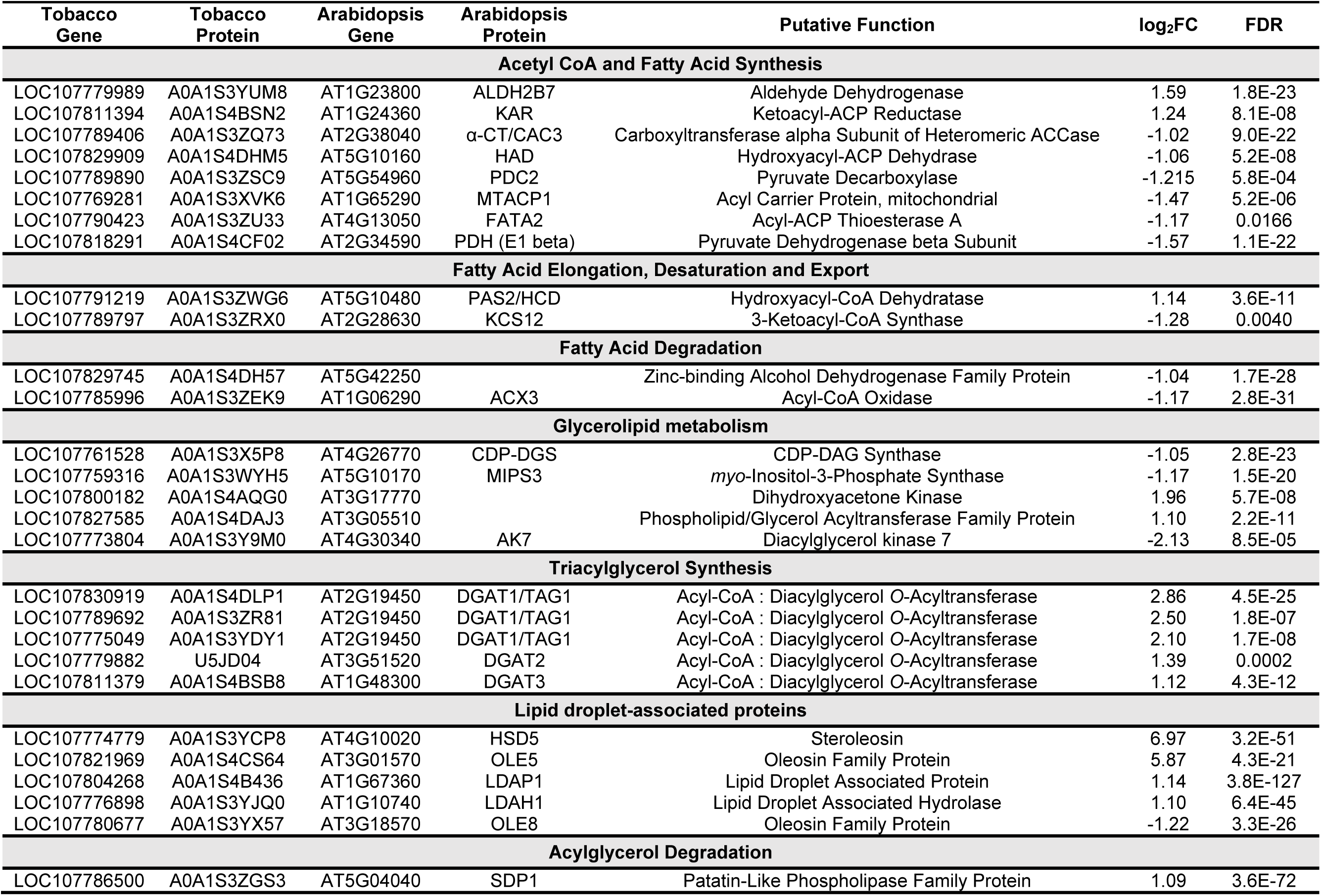
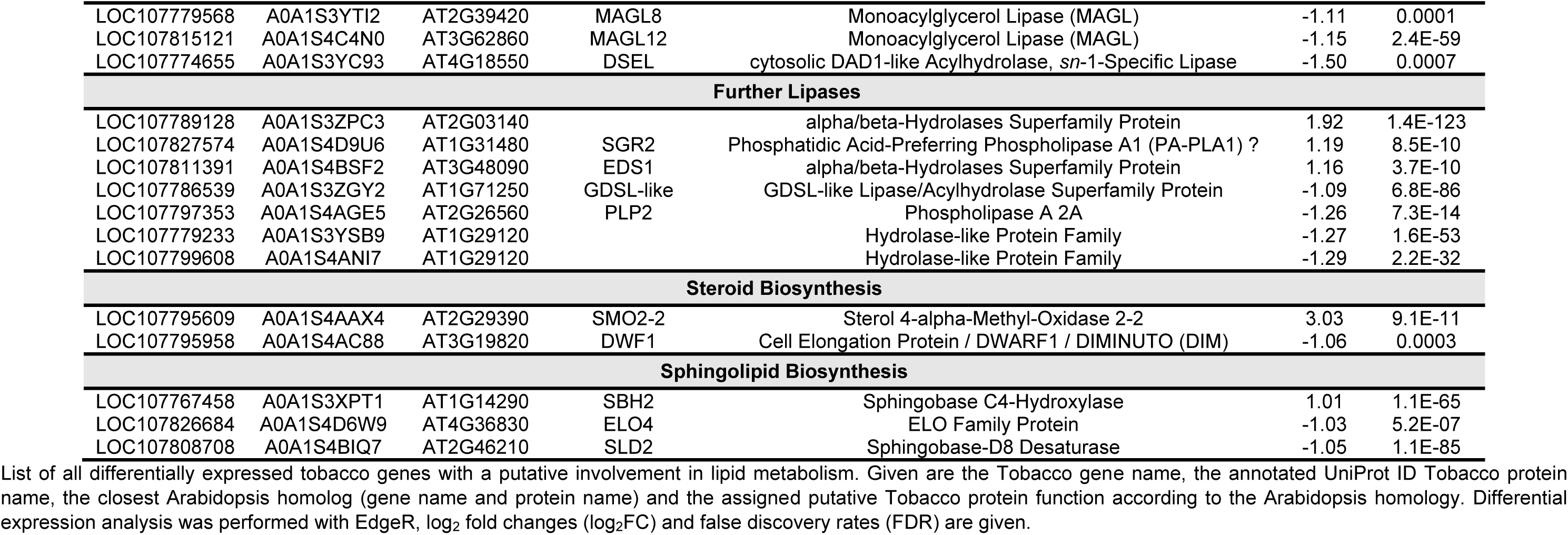
List of differentially expressed genes with putative involvement in lipid metabolism.

| Tobacco Gene | Tobacco Protein | Arabidopsis Gene | Arabidopsis Protein | Putative Function | log<sub>2</sub>FC | FDR |
| --- | --- | --- | --- | --- | --- | --- |
| **Acetyl CoA and Fatty Acid Synthesis** |  |  |  |  |  |  |
| LOC107779989 | A0A1S3YUM8 | AT1G23800 | ALDH2B7 | Aldehyde Dehydrogenase | 1.59 | 1.8E-23 |
| LOC107811394 | A0A1S4BSN2 | AT1G24360 | KAR | Ketoacyl-ACP Reductase | 1.24 | 8.1E-08 |
| LOC107789406 | A0A1S3ZQ73 | AT2G38040 | α-CT/CAC3 | Carboxyltransferase alpha Subunit of Heteromeric ACCase | -1.02 | 9.0E-22 |
| LOC107829909 | A0A1S4DHM5 | AT5G10160 | HAD | Hydroxyacyl-ACP Dehydrase | -1.06 | 5.2E-08 |
| LOC107789890 | A0A1S3ZSC9 | AT5G54960 | PDC2 | Pyruvate Decarboxylase | -1.215 | 5.8E-04 |
| LOC107769281 | A0A1S3XVK6 | AT1G65290 | MTACP1 | Acyl Carrier Protein, mitochondrial | -1.47 | 5.2E-06 |
| LOC107790423 | A0A1S3ZU33 | AT4G13050 | FATA2 | Acyl-ACP Thioesterase A | -1.17 | 0.0166 |
| LOC107818291 | A0A1S4CF02 | AT2G34590 | PDH (E1 beta) | Pyruvate Dehydrogenase beta Subunit | -1.57 | 1.1E-22 |
| **Fatty Acid Elongation, Desaturation and Export** |  |  |  |  |  |  |
| LOC107791219 | A0A1S3ZWG6 | AT5G10480 | PAS2/HCD | Hydroxyacyl-CoA Dehydratase | 1.14 | 3.6E-11 |
| LOC107789797 | A0A1S3ZRX0 | AT2G28630 | KCS12 | 3-Ketoacyl-CoA Synthase | -1.28 | 0.0040 |
| **Fatty Acid Degradation** |  |  |  |  |  |  |
| LOC107829745 | A0A1S4DH57 | AT5G42250 |  | Zinc-binding Alcohol Dehydrogenase Family Protein | -1.04 | 1.7E-28 |
| LOC107785996 | A0A1S3ZEK9 | AT1G06290 | ACX3 | Acyl-CoA Oxidase | -1.17 | 2.8E-31 |
| **Glycerolipid metabolism** |  |  |  |  |  |  |
| LOC107761528 | A0A1S3X5P8 | AT4G26770 | CDP-DGS | CDP-DAG Synthase | -1.05 | 2.8E-23 |
| LOC107759316 | A0A1S3WYH5 | AT5G10170 | MIPS3 | myo-Inositol-3-Phosphate Synthase | -1.17 | 1.5E-20 |
| LOC107800182 | A0A1S4AQG0 | AT3G17770 |  | Dihydroxyacetone Kinase | 1.96 | 5.7E-08 |
| LOC107827585 | A0A1S4DAJ3 | AT3G05510 |  | Phospholipid/Glycerol Acyltransferase Family Protein | 1.10 | 2.2E-11 |
| LOC107773804 | A0A1S3Y9M0 | AT4G30340 | AK7 | Diacylglycerol kinase 7 | -2.13 | 8.5E-05 |
| **Triacylglycerol Synthesis** |  |  |  |  |  |  |
| LOC107830919 | A0A1S4DLP1 | AT2G19450 | DGAT1/TAG1 | Acyl-CoA : Diacylglycerol O-Acyltransferase | 2.86 | 4.5E-25 |
| LOC107789692 | A0A1S3ZR81 | AT2G19450 | DGAT1/TAG1 | Acyl-CoA : Diacylglycerol O-Acyltransferase | 2.50 | 1.8E-07 |
| LOC107775049 | A0A1S3YDY1 | AT2G19450 | DGAT1/TAG1 | Acyl-CoA : Diacylglycerol O-Acyltransferase | 2.10 | 1.7E-08 |
| LOC107779882 | U5JD04 | AT3G51520 | DGAT2 | Acyl-CoA : Diacylglycerol O-Acyltransferase | 1.39 | 0.0002 |
| LOC107811379 | A0A1S4BSB8 | AT1G48300 | DGAT3 | Acyl-CoA : Diacylglycerol O-Acyltransferase | 1.12 | 4.3E-12 |
| **Lipid droplet-associated proteins** |  |  |  |  |  |  |
| LOC107774779 | A0A1S3YCP8 | AT4G10020 | HSD5 | Steroleosin | 6.97 | 3.2E-51 |
| LOC107821969 | A0A1S4CS64 | AT3G01570 | OLE5 | Oleosin Family Protein | 5.87 | 4.3E-21 |
| LOC107804268 | A0A1S4B436 | AT1G67360 | LDAP1 | Lipid Droplet Associated Protein | 1.14 | 3.8E-127 |
| LOC107776898 | A0A1S3YJQ0 | AT1G10740 | LDAH1 | Lipid Droplet Associated Hydrolase | 1.10 | 6.4E-45 |
| LOC107780677 | A0A1S3YX57 | AT3G18570 | OLE8 | Oleosin Family Protein | -1.22 | 3.3E-26 |
| **Acylglycerol Degradation** |  |  |  |  |  |  |
| LOC107786500 | A0A1S3ZGS3 | AT5G04040 | SDP1 | Patatin-Like Phospholipase Family Protein | 1.09 | 3.6E-72 |
| LOC107779568 | A0A1S3YTI2 | AT2G39420 | MAGL8 | Monoacylglycerol Lipase (MAGL) | -1.11 | 0.0001 |
| LOC107815121 | A0A1S4C4N0 | AT3G62860 | MAGL12 | Monoacylglycerol Lipase (MAGL) | -1.15 | 2.4E-59 |
| LOC107774655 | A0A1S3YC93 | AT4G18550 | DSEL | cytosolic DAD1-like Acylhydrolase, *sn*-1-Specific Lipase | -1.50 | 0.0007 |
| **Further Lipases** |  |  |  |  |  |  |
| LOC107789128 | A0A1S3ZPC3 | AT2G03140 |  | alpha/beta-Hydrolases Superfamily Protein | 1.92 | 1.4E-123 |
| LOC107827574 | A0A1S4D9U6 | AT1G31480 | SGR2 | Phosphatidic Acid-Preferring Phospholipase A1 (PA-PLA1) ? | 1.19 | 8.5E-10 |
| LOC107811391 | A0A1S4BSF2 | AT3G48090 | EDS1 | alpha/beta-Hydrolases Superfamily Protein | 1.16 | 3.7E-10 |
| LOC107786539 | A0A1S3ZGY2 | AT1G71250 | GDSL-like | GDSL-like Lipase/Acylhydrolase Superfamily Protein | -1.09 | 6.8E-86 |
| LOC107797353 | A0A1S4AGE5 | AT2G26560 | PLP2 | Phospholipase A 2A | -1.26 | 7.3E-14 |
| LOC107779233 | A0A1S3YSB9 | AT1G29120 |  | Hydrolase-like Protein Family | -1.27 | 1.6E-53 |
| LOC107799608 | A0A1S4ANI7 | AT1G29120 |  | Hydrolase-like Protein Family | -1.29 | 2.2E-32 |
| **Steroid Biosynthesis** |  |  |  |  |  |  |
| LOC107795609 | A0A1S4AAX4 | AT2G29390 | SMO2-2 | Sterol 4-alpha-Methyl-Oxidase 2-2 | 3.03 | 9.1E-11 |
| LOC107795958 | A0A1S4AC88 | AT3G19820 | DWF1 | Cell Elongation Protein / DWARF1 / DIMINUTO (DIM) | -1.06 | 0.0003 |
| **Sphingolipid Biosynthesis** |  |  |  |  |  |  |
| LOC107767458 | A0A1S3XPT1 | AT1G14290 | SBH2 | Sphingobase C4-Hydroxylase | 1.01 | 1.1E-65 |
| LOC107826684 | A0A1S4D6W9 | AT4G36830 | ELO4 | ELO Family Protein | -1.03 | 5.2E-07 |
| LOC107808708 | A0A1S4BIQ7 | AT2G46210 | SLD2 | Sphingobase-D8 Desaturase | -1.05 | 1.1E-85 |

List of all differentially expressed tobacco genes with a putative involvement in lipid metabolism. Given are the Tobacco gene name, the annotated UniProt ID Tobacco protein name, the closest Arabidopsis homolog (gene name and protein name) and the assigned putative Tobacco protein function according to the Arabidopsis homology. Differential expression analysis was performed with EdgeR, log<sub>2</sub> fold changes (log<sub>2</sub>FC) and false discovery rates (FDR) are given.

This result already indicates that lipid adaptations are not strongly reflected on a transcriptional level. To take a closer look at the differentially expressed genes, lipid genes were grouped according to their pathways.

### Fatty acid synthesis is not transcriptionally upregulated

Pollen tube growth requires fatty acid synthesis in the plastids to cope with the increasing need for membrane lipids in growing pollen tubes (Ischebeck, 2016). Under HS, pollen tubes seem to need slightly more membrane lipids, as several minor lipid classes as well as TG levels accumulated stronger at high temperature (Figure 2A, 5B). However, neither of the pathways leading to acetyl-CoA synthesis via the pyruvate dehydrogenase or pyruvate decarboxylase seem to be strongly affected (Supplemental Dataset S24). The same is true for fatty acid synthesis itself, as only one relevant gene, one of six putative ketoacyl-ACP reductases (KAR) was upregulated It was previously shown that downregulation of the ketoacylsynthase KASII/FAB1, which is responsible for the elongation from C16 to C18 acyl chains, leads to strongly increased levels of C16 acyl chains in Arabidopsis (Pidkowich *et al.*, 2007). Interestingly, none of the four isoforms was considerably downregulated in tobacco pollen tubes under HS, despite the increase of 16:0 acyl chains across lipid classes (Figure 3).

What is more, no differentially expressed fatty acid desaturases (FADs) were detected in our screen. FAD2 and FAD3, the desaturases that catalyse in PC the desaturation from 18:1 to 18:2 and from 18:2 to 18:3, respectively, are not downregulated but several isoforms were slightly upregulated.

### Transcript does not reflect adaptations in glycerolipid profiles

The expression levels of enzymes involved in glycerolipid metabolism gave an indifferent picture. While some individual isoforms were differentially expressed (Table 4), only few clear trends could be observed. Two putative PA phosphatases homologous to Arabidopsis LPP1/PAP1 had very high abundance levels and increased by 25 % and 55 % during HS, respectively. Also two of three CGI58-type lysophosphatidic acid acyltransferases that are involved in membrane and neutral lipid homeostasis (Ghosh *et al.*, 2009; James *et al.*, 2010) were slightly upregulated by 32 % and 35 %, respectively (Supplementary dataset S19). Likewise, no clear trends for genes involved in galactoglycerolipid metabolism were observed.

Transcripts important for sulfolipid metabolism were not detected, indicating again that these lipids do not occur in tobacco pollen tubes.

### Transcripts hint at a dynamic turnover of TG upon heat stress

Taking a closer look at TG metabolism, we found that diacylglycerol acyltransferase 1 (DGAT1), DGAT2 and DGAT3 homologues are strongly transcriptionally upregulated after 3 h of HS. However, a homologue of the major TG lipase SUGAR DEPENDENT 1 (SDP1) is also upregulated, so is another putative TG lipase. Upregulated genes also include some known LD-localised proteins, a putative 11-β−hydroxysteroid dehydrogenase (HSD) is very strongly upregulated (125-fold), which is interesting as previously no HSDs were found on LDs of tobacco pollen tubes on the protein level (Kretzschmar *et al.*, 2018). Also, an oleosin, a lipid droplet-associated protein (LDAP) and a putative lipid droplet-associated hydrolase (LDAH) (Kretzschmar *et al.*, 2020) show upregulation (Table 4); while another putative oleosin family protein is downregulated. Some low but significant changes can be observed in the scaffold protein plant UBX-domain containing protein (PUX10), that plays a role in the degradation of LD proteins (Deruyffelaere *et al.*, 2018; Kretzschmar *et al.*, 2018). All four detected putative homologues show slight upregulation (Supplemental Dataset S24).

### Several genes involved in sterol and sphingolipid metabolism show differential expression

A transcript encoding for a methylsterol monooxygenase 2 (SMO2)-like isoform showed 8-fold upregulation. SMO proteins are involved in sterol synthesis, to be more precise SMO2-1 and SMO2-2 are involved in the reaction from 24-methylenelophenol to episterol (precursor of campesterol, brassinosteroids and brassicasterol) and from 24-ehtylenelophenol to δ7-avenasterol (that can be converted to isofucosterol, sitosterol and stigmasterol later). Furthermore, a putative δ(24)-sterol reductase shows two-fold upregulation.

Regarding sphingolipid metabolism, a Δ8-fatty-acid desaturase-like protein that shares 62 % sequence identity with the Arabidopsis SPHINGOID LONG CHAIN BASE DESATURASE 2 (SLD2) was two-fold downregulated on transcript level, while other homologues of this enzyme showed upregulation (Table 4, Supplemental Dataset S24). The transcript of a putative sphingoid base hydroxylase 2 (SBH2) that inserts a hydroxyl group at the C4 position showed two-fold upregulation. One of three isoforms of dihydrosphingosine Δ-4 desaturases catalysing a competing reaction, the insertion of a double bond at the Δ4 position, was downregulated by 31 %, while the other isoforms displayed little change.

### Differentially expressed genes suggest involvement of hormones in heat stress adaptation

Some other interesting differentially expressed genes that have not been covered so far are highlighted in Table 5. The gene displaying the strongest upregulation is annotated as “uncharacterised protein” in tobacco and its closest Arabidopsis homologue, AT3G10020, is annotated as “plant/protein”. Publications suggest that it is a stress-responsive gene. Strongest downregulation was observed for a homologue of BON association protein 2 (BAP2), a reported inhibitor of programmed cell death (Yang *et al.*, 2007).

**Table 5:** Selection of other differentially expressed genes of interest.

| Tobacco Gene | Tobacco Protein | Arabidopsis Gene | Arabidopsis Protein | Putative Function | log<sub>2</sub>FC | log<sub>2</sub>CPM | FDR |
| --- | --- | --- | --- | --- | --- | --- | --- |
| LOC107766703 | A0A1S3XM78 | AT3G10020 | unknown protein | plant/protein | 12.07 | 3.38 | 0 |
| LOC107778746 | A0A1S3YRE7 | AT5G05600 | JAO2 | 2-oxoglutarate (2OG) and Fe(II)-dependent oxygenase superfamily protein | 10.23 | 3.19 | 0 |
| LOC107817403 | A0A1S4CCK8 | AT3G22370 | AOX1a | alternative oxidase | 9.05 | 0.41 | 9.24E-59 |
| LOC107759427 | A0A1S3WZ93 | AT3G10020 | unknown protein | plant/protein | 7.89 | 2.57 | 2.52E-241 |
| LOC107804414 | A0A1S4B4D1 | AT5G03370 |  | acylphosphatase family | 6.96 | -1.40 | 2.73E-15 |
| LOC107774730 | A0A1S3YCX4 | AT3G09640 | APX2 | ascorbate peroxidase | 5.29 | -1.31 | 3.20E-14 |
| LOC107788327 | A0A1S3ZM96 | AT3G47520 | MDH | malate dehydrogenase | 4.07 | -0.51 | 5.52E-20 |
| LOC107795525 | A0A1S4AAK3 | AT1G14130 | DAO1 | 2-oxoglutarate (2OG) and Fe(II)-dependent oxygenase superfamily protein | 3.88 | 2.70 | 4.11E-184 |
| LOC107801843 | A0A1S4AVW8 | AT5G47530 |  | Auxin-responsive family protein | 3.44 | -0.79 | 2.59E-14 |
| LOC107769851 | A0A1S3XY09 | AT3G21865 | PEX22 | peroxin | 3.40 | 0.28 | 4.42E-27 |
| LOC107764611 | A0A1S3XFN6 | AT5G53120 | SPDS3 | spermidine synthase | 3.15 | 2.26 | 6.03E-95 |
| LOC107808988 | A0A1S4BJJ5 | AT5G24240 | ATPI4KGAMMA3 | phosphatidylinositol 4-kinase gamma-like protein | 2.97 | 3.59 | 1.35E-219 |
| LOC107817192 | A0A1S4CB63 | AT4G27510 |  | 2-isopropylmalate synthase | 2.36 | -0.94 | 1.05E-07 |
| LOC107762352 | A0A1S3X8G8 | AT1G49340 | ATPI4KALPHA | Phosphatidylinositol 3- and 4-kinase family protein | 2.36 | 1.92 | 7.99E-50 |
| LOC107770164 | A0A075EYT2 | AT1G17710 | PEPC1 | phosphoethanolamine/phosphocholine phosphatase | 2.34 | 3.66 | 1.55E-173 |
| LOC107808981 | A0A1S4BJF1 | AT1G17710 | PEPC1 | phosphoethanolamine/phosphocholine phosphatase | 2.00 | 5.00 | 9.42E-139 |
| LOC107817132 | A0A1S4CB76 | AT1G17710 | PEPC1 | phosphoethanolamine/phosphocholine phosphatase | 1.50 | 2.74 | 3.69E-38 |
| LOC107799822 | A0A1S4APJ5 | AT2G43020 | PAO2 | polyamine oxidase | 1.17 | 4.50 | 2.81E-72 |
| LOC107768947 | A0A1S3XUI8 | AT1G65930 | cICDH | cytosolic NADP+-dependent isocitrate dehydrogenase | -1.01 | 3.04 | 6.39E-13 |
| LOC107828000 | A0A1S4DBY9 | AT2G28350 | ARF10 | auxin response factor | -1.46 | 0.28 | 1.39E-07 |
| LOC107767522 | A0A1S3XQ16 | AT4G33430 | BAK1 | BRI1-associated receptor kinase | -1.47 | -0.85 | 0.00091587 |
| LOC107798522 | T1WMC3 | AT1G30135 | JAZ8 | jasmonate-zim-domain protein | -1.49 | -0.40 | 0.00010296 |
| LOC107759371 | A0A1S3WYR7 | AT4G34710 | ADC2 | arginine decarboxylase | -1.54 | 6.27 | 4.78E-241 |
| LOC107819571 | A0A1S4CJ33 | AT1G02170 | MC1 | metacaspase | -2.00 | -1.31 | 0.00123282 |
| LOC107832546 | A0A1S4DR37 | AT5G25260 | FLOT2 | Flotillin | -2.07 | -0.44 | 1.32E-06 |
| LOC107820452 | A0A1S4CLU4 | AT1G02170 | MC1 | metacaspase | -2.57 | 1.00 | 7.54E-20 |
| LOC107820447 | A0A1S4CLX5 | AT1G02170 | MC1 | metacaspase | -2.72 | -0.41 | 1.84E-09 |
| LOC107805149 | A0A1S4B6Y7 | AT5G04200 | MC9 | metacaspase | -2.86 | -1.31 | 1.93E-06 |
| LOC107824234 | A0A1S4CZ94 | AT2G19800 | MIOX2 | myo-inositol oxygenase | -2.89 | -0.54 | 2.29E-09 |
| LOC107773293 | A0A1S3Y7Q9 | AT2G19800 | MIOX2 | myo-inositol oxygenase | -3.17 | 0.20 | 1.86E-15 |
| LOC107799833 | A0A1S4APP9 | AT2G45760 | BAP2 | BON association protein | -4.32 | -0.46 | 7.69E-16 |

Selection of differentially expressed Tobacco genes (including the gene with the strongest up- and downregulation, respectively), the annotated UniProt ID Tobacco protein name, the closest Arabidopsis homolog (gene name and protein name) and the assigned putative Tobacco protein function according to the Arabidopsis homology. Differential expression analysis was performed with EdgeR, log<sub>2</sub> fold changes (log<sub>2</sub>FC), log<sub>2</sub> counts per million (log<sub>2</sub>CPM) and false discovery rates (FDR) are given.

While auxins were already mentioned in the GO-terms, some other hormones might also play a role in HS adaptation of tobacco pollen tubes. Strong upregulation was observed for a homologue of Arabidopsis JASMONATE-INDUCED OXYGENASE2 (JAO2), annotated as a protein with similarity to flavonol synthases and involved in the detoxification of polycyclic aromatic hydrocarbons (Hernández-Vega *et al.*, 2017). JASMONATE-ZIM-DOMAIN PROTEIN 8 (JAZ8) on the other hand was transcriptionally downregulated. Transcript of a putative homologue of BRI1- associated receptor kinase 1 (BAK1), a receptor-like kinase in brassinosteroid sensing, was downregulated under HS.

## Discussion

### The tobacco pollen tube transcriptome shows expected but also unique adaptations to heat stress

As shown in several previous studies on plants, HS leads to a rapid and strong remodelling of the transcriptome (Kotak *et al.*, 2007; Mittal, Madhyastha and Grover, 2012; Rahmati Ishka *et al.*, 2018). One reaction conserved across pro- and eukaryotes is the upregulation of heat shock proteins (HSPs) (Jacob, Hirt and Bendahmane, 2017). Accordingly, most transcripts strongly upregulated in response to HS encode HSPs in this study as well indicating that the applied temperature regime a imposes heat stress to the pollen tubes (Supplementary dataset S16). This upregulation has also been found to be a key response under HS in developing pollen, prior to pollen tube growth (Fragkostefanakis *et al.*, 2016; Keller *et al.*, 2018). However, we also found differences between HS responses in tobacco pollen tubes and vegetative tissues of Arabidopsis, as several homologs of heat-induced genes in Arabidopsis are expressed, but not upregulated upon HS in tobacco pollen tubes. These include genes encoding for LATE EMBRYOGENESIS ABUNDANT (LEA) proteins, which protect other proteins and membranes especially during desiccation but are also upregulated by HS (Priya *et al.*, 2019). On the contrary, we found genes upregulated that have so far not been associated with HS. These include for example a Flotillin-like gene with homology to Arabidopsis FLOT2 (Table 5). Flotillins are involved in formation of membrane microdomains (Haney and Long, 2010; Li *et al.*, 2012). These data suggest, that Flotillins might also play an important role during pollen tube growth, maybe especially so under stress. The presence of sterol-rich membrane microdomains containing flotillin-like protein in rice pollen has recently been shown (Han *et al.*, 2018).

### Heat-induced glycerolipid remodelling does not appear to be strongly transcriptionally controlled

The expression data shows no conclusive picture as to how the membrane lipid composition is altered. Nevertheless, several conclusions can be drawn from membrane lipid remodelling: Especially striking is the increase of the saturated acyl chains 16:0 and 18:0, while the ratio of monounsaturated to polyunsaturated acyl chains is not severely altered. This result suggests a regulation of fatty acid synthesis in the plastids, where it is determined if the acyl carrier protein (ACP)-connected acyl chains are initially elongated and desaturated. Only after elongation and desaturation from 16:0 to 18:1, the acyl chains can be precursors for further extraplastidial desaturation.

Arabidopsis and likely also other plant species harbour three types of ketoacylsynthases (KAS) with only one, KASII/FAB1, being responsible for the elongation of 16:0-ACP to 18:0-ACP (Wu *et al.*, 1994; Pidkowich *et al.*, 2007). KASII/FAB1 would thus pose a good target for regulation. Regulation of the Δ9 desaturase FAB2 could be a further factor, as it introduces the first double bond to form 18:1. Furthermore, thioesterases might play a role, as knockout of FATB leads to a strong reduction of saturated acyl chains (Bonaventure *et al.*, 2003), and knockdown of the two FATA genes also influences acyl composition (Moreno-Pérez *et al.*, 2012).

One can speculate that during HS the *de novo* synthesised fatty acids are even stronger saturated than reflected by the membrane lipid composition, as some acyl chains were likely already synthesised prior to HS. This would imply a rapid and strong adaptation of the above mentioned enzymes, likely through protein degradation, post-translational modification or a direct influence of temperature on the enzymatic activities. A transcriptional regulation is however unlikely, as respective transcripts were little or not changed (Supplemental Dataset S24).

### Role of phosphatidylserine and galactoglycerolipids in heat stress adaptation

Some lipid classes increase comparably stronger under HS. This is especially true for PS and DGDG. PS is found in the cytosolic leaflets of the plasma membrane and in endosomes and is negatively charged. It can form nanodomains in the plasma membrane that are important for Rho signalling (Platre *et al.*, 2019). These small G proteins have also been shown to be important for the regulation of pollen tube growth (Scholz *et al.*, 2020), but it remains to be studied if they also depend on PS, how PS is distributed in the pollen tube, and if this distribution changes under HS. Galactoglycerolipids are the most abundant lipids of thylakoids and there is strong galactoglycerolipid remodeling under temperature stress in Arabidopsis (Chen *et al.*, 2006; Higashi *et al.*, 2015) and tomato leaves (Spicher, Glauser and Kessler, 2016). In agreement with studies concerning other class of membranes lipids, the level of unsaturation in galactoglycerolipids is inversely correlated with growth temperature. Furthermore, heat acclimation at 38 °C in wild-type Arabidopsis increases DGDG, DGDG to MGDG ratio, and the saturation level of DGDG (Higashi *et al.*, 2015). Also, a role for galactoglycerolipids in acquired thermotolerance was already described in a study in 2006, when the authors did a mutant screen for plants defective in the acquisition of thermotolerance and found a mutant of *DGD1* (Chen *et al.*, 2006). While pollen tubes contain plastids, these harbour no thylakoids (Staff *et al.*, 1989) and galactoglycerolipids in pollen tubes were discussed to be especially important in extraplastidial membranes, where they are also formed in phosphate-starved vegetative tissues (Härtel, Dörmann and Benning, 2000). Evidence for the abundance of galactoglycerolipids in the male gametophyte comes from glycerolipid profiling of lily pollen tubes before and after elongation revealing a 5.7-fold increase in DGDG and a 2.8-fold increase in the MGDG levels (Nakamura, Kobayashi and Ohta, 2009). The use of the MGDG synthase inhibitor galvestine-1 further highlighted an important role of galactoglycerolipids in pollen tube growth and the use of a specific antibody indicated a DGDG localization at the periphery of Arabidopsis pollen tubes most probably at the plasma membrane (Botté *et al.*, 2011). In addition, the fact that galactoglycerolipids account for 11 % of all membrane forming glycerolipids in tobacco pollen tubes (Müller and Ischebeck, 2018) speaks for their presence in extraplastidial membranes.

### Transcript and lipidome suggest dynamic adaptations in LD-turnover and TG metabolism

While transcripts involved in lipid metabolism were mostly unaffected by HS, transcripts coding for genes involved in TG turnover and LD biology showed more dynamic adaptations to HS (Supplemental Dataset S24). An increase of TG levels under HS has already been observed, e.g. in the algal species *Nannochloropsis oculata* (Converti *et al.*, 2009), *Ettlia oleoabundans* (Yang *et al.*, 2013), *Coccomyxa subellipsoidea* C169 (Allen *et al.*, 2018), or *Chlamydomonas reinhardtii* (Légeret *et al.*, 2016), but also in Arabidopsis leaves (Higashi *et al.*, 2015; Shiva *et al.*, 2020) and seedlings (Mueller *et al.*, 2015), and tomato fruits (Almeida, Perez-Fons and Fraser, 2021). In Arabidopsis seedlings, PHOSPHOLIPID:DIACYLGLYCEROL ACYLTRANSFERASE1 (PDAT1), an enzyme transferring a fatty acid from PC to DG to yield TG, is necessary for heat-induced TG accumulation. Also, the *pdat1* mutant seedlings were more sensitive to HS, indicating that PDAT1-mediated TG accumulation mediates thermotolerance (Mueller *et al.*, 2017). Another interesting fact hinting at an involvement of not just TG but LDs as organelles in HS adaptations is the 125-fold upregulation of a putative HSD5 homologue (Table 4) suggesting for an involvement of LDs in HS adaptation beyond TG accumulation. HSDs, also called steroleosins, are important for plant development and are involved in stress responses as well as wax metabolism (Li *et al.*, 2007; Zhang *et al.*, 2016; Shao *et al.*, 2019). Steroleosins are presumably involved in brassinosteroid metabolism (Li *et al.*, 2007) but were previously not found on LDs of tobacco pollen tubes (Kretzschmar *et al.*, 2018). In this study, only two transcripts with very low expression levels were detected at non-stressed conditions.

### Changes in sterol *lipid* and sphingolipid metabolism might contribute to heat adaption

In contrast to TG that increases during HS and might therefore act as a sink for acyl chains, SE species are decreased, indicating that free sterols are released during heat adaptation and could help to adapt membrane fluidity (Dufourc, 2008). Free sterols could furthermore be converted to SG and ASG species, which were shown to increase upon HS. As so far, no plant sterol esterase has been described (Ischebeck *et al.,* 2020), it is not clear if SE breakdown is regulated transcriptionally.

Alterations in sphingolipid composition on the other hand could at least in part be explained by changes in transcript levels – the slight increase of hydroxylation versus desaturation at the C4 position of the SPB in HexCers is in line with the respective changes in the transcripts coding for the responsible enzymes (Table 4; Supplemental Dataset S24). The increased hydroxylation of the SPB moiety in HexCers and the acyl chain in Hex-HexNAc-GlcA-IPCs could stabilise sphingolipid-containing microdomains through their ability to form lateral hydrogen bonds with other sphingolipids or free sterols (Mamode Cassim *et al*., 2019).

The strong increase of SPBP on the other hand is not reflected on the transcript level as none of most abundant SPB kinases are strongly upregulated (Supplemental Dataset S24). The production of SPBP is thought to play a role in balancing the levels of putatively toxic SPBs. Furthermore, SPB breakdown requires its phosphorylation to SPBP followed by degradation to PE and palmitic aldehyde by SPBP lyase. Furthermore, SPBP could have signalling function (Puli *et al.*, 2016).

### Metabolomic adaptations might facilitate thermotolerance

In previous studies on heat-stressed shoot tissues of Arabidopsis (Kaplan *et al.*, 2004; Harsh *et al.*, 2016; Zinta *et al.*, 2018; Lawas *et al.*, 2019), it was found that different sugars accumulated, including glucose, fructose and in some cases sucrose. In tobacco pollen tubes, the heat-induced accumulation of sugars was rather modest in comparison and the somewhat higher increase in sucrose has to be considered with care, as sucrose was also contained in the growth medium (Figure 9). Interestingly though, we found a strong increase in sedoheptulose, a seven carbon sugar normally occurring in its phosphorylated form in the Calvin-Benson cycle and the pentose phosphate pathway. It is unclear if this sugar has any protective function, but it was also found to highly accumulate in the alga *Phaeodactylum tricornutum* under nitrogen deprivation (Popko *et al.*, 2016).

In addition, an increase of several free amino acids was observed in under heat stress (Figure 9). Proline levels, however, stayed relatively stable and although it accumulates very strongly after 9 h of pollen tube growth, it does so under normal as well as stress conditions. This is interesting, as proline was found to be clearly heat-inducible in studies on shoot tissues (Harsh *et al.*, 2016; Zinta *et al.*, 2018; Lawas *et al.*, 2019). However, a study on potato leaves showed that HS alone does not have significant influence on proline levels, neither in stress-susceptible nor in resistant potato cultivars. In the study, proline only accumulated after drought or combined HS and drought stress (Demirel *et al.*, 2020). The same study observed an increase in lysine in one of the stress-susceptible cultivars following HS. It is possible that pipecolate, similar in structure to proline, plays a role in adaptation to heat in pollen tubes, as pipecolate was more heat-responsive than proline in the present study. Pipecolate was also shown to strongly accumulate during tobacco pollen development (Rotsch *et al.*, 2017) and it increases in shoot tissues upon pathogen infection (Návarová *et al.*, 2013; Ding *et al.*, 2016).

## Conclusion

Adaptation of tobacco pollen tubes to heat stress appears to be a complex and multi-layered trait that requires intricate interplay of different cellular processes. A rapid lipid remodelling that is not controlled transcriptionally is equally as important as metabolomic adaptations and transcriptional changes. Therefore, the present study with several large datasets poses a useful resource for future studies on heat stress tolerance.

## Materials and Methods

### Plant material and growth conditions

Tobacco (*Nicotiana tabacum* L. cv. Samsun-NN) plants were grown in the greenhouse as previously described (Rotsch *et al.*, 2017): plants were kept under 14 h of light from mercury-vapor lamps in addition to sunlight with light intensities of 150 – 300 µmol m^−2^ s^−1^ at flowers and 50 – 100 µmol m^−2^ s^−1^ at mid-height leaves.

Temperature was set to 16 °C at night and 21 °C during the day with a relative humidity of 57–68 %.

Anthers were harvested from flower buds right before anthesis and dried at RT for up to four days before pollen from the anthers was collected by sieving. Pollen were weighed and rehydrated for 10 min in liquid pollen tube medium (5 % w/v sucrose, 12.5 % w/v PEG-4000, 15 mM MES-KOH pH 5.9, 1 mM CaCl_2_,1 mM KCl, 0.8 mM MgSO_4_, 0.01 % H_3_BO_3_, 30 µM CuSO_4_) modified from (Read, Clarke and Bacic, 1993). The indicated amounts were spread onto cellophane foil (Max Bringmann KG, Germany) and placed on 50 ml of solid pollen tube medium (2 % Agarose, 5 % sucrose, 6 % PEG-4000, 15 mM MES-KOH pH 5.9, 1 mM CaCl_2_,1 mM KCl, 0.8 mM MgSO_4_, 0.01 % H_3_BO_3_, 30 µM CuSO_4_) inside square petri dishes (120 mm x 120 mm x 17 mm with vents, Greiner Bio-One, Kremsmünster, Austria), sealed with MicroporeTM (3M, Saint Paul, MN, USA). All pollen tubes were grown for 3 h at RT. Control pollen tubes were kept at RT for another 3 h, while heat stressed-pollen tubes were transferred to 37 °C for 3 h. For HSR, pollen tubes were then transferred to RT again for the next 3 h, while non-relieved pollen tubes remained at 37 °C. Pollen tubes were harvested using spatula and forceps, and directly transferred into extraction solution for lipid or metabolite analysis. For RNA extraction the material was immediately flash-frozen for further processing.

### Lipidomic analysis

For lipidomic analyses with an ultra-performance liquid chromatography (UPLC) system coupled with a chip-based nanoelectrospray ionization (nanoESI) source and a triple quadrupole analyzer for tandem mass spectrometry (MS/MS) (Herrfurth, Liu and Feussner, 2021), 20 mg of freeze-dried pollen tube tissue were ground and extracted with 60:26:14 v/v/v propan-2-ol/hexane/water according to Markham *et al.*, 2006. After extraction, the sample was dissolved in 0.8 ml 4:4:1 v/v/v tetrahydrofuran/methanol/water. UPLC-nanoESI-MS/MS analysis of molecular species from 36 different lipid classes was performed with lipid subclass-specific parameters as shown in Supplemental Table S1. For reverse-phase LC separation, an ACQUITY UPLC^®^ I-class system (Waters Corp., Milford, MA, USA) equipped with an ACQUITY UPLC^®^ HSS T3 column (100 mm × 1 mm, 1 μm; Waters Corp., Milford, MA, USA) was used. Solvent A was 3:7 v/v methanol/20mM ammonium acetate containing 0.1 % v/v acetic acid, and solvent B was 6:3:1 v/v/v tetrahydrofuran/methanol/20 mM ammonium acetate containing 0.1 % v/v acetic acid. All lipid subclasses were separated with linear binary gradients following the same scheme: respective start condition (Supplemental Table S1) held for 2 min, linear increase to 100 % solvent B for 8 min, 100 % solvent B held for 2 min and re-equilibration to start conditions in 4 min. Chip-based nanoESI was achieved with a TriVersa Nanomate^®^ (Advion, Ithaca, NY, USA) in the positive or negative ion mode, respectively (Supplemental Table S1). Lipid molecular species were detected with a 6500 QTRAP^®^ tandem mass spectrometer (AB Sciex, Framingham, MA, USA) by multiple reaction monitoring (MRM) with lipid class-specific parameters (Supplemental Table S1). For glycerolipid and glycerophospholipid analysis, all mass transitions for the distinct lipid subclasses were measured for the putative lipid species having 16:0, 16:1, 16:2, 16:3, 17:0, 17:1, 17:2, 17:3, 18:0, 18:1, 18:2, 18:3, 19:0, 19:1, 19:2, 19:3, 20:0, 20:1, 20:2, 22:0, 22:1, 24:0, 24:1, 26:0, and 26:1 as acyl residues. For sphingolipid analysis, all mass transitions for the distinct lipid subclasses were measured for the putative lipid species having 18:0;O2, 18:1;O2, 18:2;O2, 18:0;O3, and 18:1;O2 as SPB residues and chain lengths from C16 to C28 as fatty acid residues that are saturated or monounsaturated and unhydroxylated or monohydroxylated. For sterol lipid analysis, all mass transitions for the distinct classes were measured for the putative lipid species having cholesterol, sitosterol, campesterol, stigmasterol/putative methylenepollinastanol/isofucosterol/24- methylenelophenol, 24-methylenecholesterol/ Δ5,24-ergostadienol, and cycloeucalenol as steryl residue and the fatty acids as acyl residues as described above for the glycerolipid and glycerophospholipid analysis. To improve the chromatographic separation and mass spectrometric detection of distinct lipid subclasses, phosphate groups and hydroxyl groups of the respective lipid compounds were chemically modified by methylation (LPA, PA, PIP and PIP_2_) modified from Lee *et al*., 2013 or by acetylation (CerP and SPBP) modified from Berdyshev *et al.* 2005, respectively. The calculation of the relative abundance of all detected lipid subclasses and of relative lipid subclass profiles is described in the Supplemental Datasets S1-S6 and S9-S11. All calculations were made according to the following procedure.

1. Correction of the absolute peak area with the naturally occurring proportion of the ^13^C isotopes (isotopic correction factor, icf);
2. Addition of icf-corrected absolute peak area from corresponding mass transitions;
3. Total peak area of a lipid subclass (addition of all detected molecular species of the respective lipid subclass) and normalization to the values after 3 h of RT growth (relative abundance compared to RT 3h); relative proportion of the lipid species-specific peak area on the total peak area of the respective lipid subclass (relative lipid subclass profile).

### Lipid extraction and fatty acid methyl esterification

For analyses by gas chromatography flame ionisation detection (GC-FID), total lipids from pollen tubes grown from 20 mg of dry pollen were extracted by methyl-*tert*-butyl ether (MTBE) extraction (modified from Matyash *et al.*, 2008). Pollen tubes were harvested into 2 mL of pre-heated propan-2-ol (75 °C) and kept at 75 °C for 5-10 min.

0.05 mg triheptadecanoate (Merck KGaA, Darmstadt, Germany) in 50 µL chloroform as internal standard for absolute quantification was directly added to the propan-2-ol. Propan-2-ol was transferred to a new vial and evaporated under N_2_stream, the pollen tubes were covered with 2 mL of 3:1 v/v MTBE/methanol. Pollen tube tissue was disrupted with a spatula, vortexed and then shaken at 4 °C for one hour. After 5 minutes of centrifugation at 1000 x g, the supernatant was transferred to the vial with the evaporated propan-2-ol and 1 mL of 0.9 % w/v NaOH was added. Samples were vortexed and centrifuged again before the upper phase was transferred to a new vial. Solvent was evaporated under N_2_ stream and samples dissolved in 250 µL 65:56:8 v/v/v chloroform/methanol/water.

The total lipids were then separated by thin layer chromatography (TLC). 80 µL of the dissolved samples were spotted with a TLC spotter to TLC plates (TLC Silica gel 60, Merck KGaA). For extraction of TGs, plates were run in 80:20:1 v/v/v hexane/diethylether/acetic acid. The bands comigrating with the TG standard were scratched out. To obtain fatty acid methyl esters (FAMEs), samples were then subjected to 1 mL FAME reagent (2.5 % v/v H_2_SO_4_, 2 % v/v dimethoxypropane in 2:1 v/v methanol/toluene and under constant shaking incubated at 80 °C in a water bath for one hour. The reaction was stopped by adding 1 mL of saturated NaCl solution and vortexing. FAMEs were then extracted adding 1 mL of hexane, centrifuging 10 minutes at 2,000 x g and transferring the upper phase to a new vial. Hexane was evaporated and samples resuspended in 25 µL of acetonitrile for subsequent GC-FID analysis.

### Central metabolite and sterol extraction and derivatisation

Primary metabolite and sterol extraction were performed as previously described (Rotsch *et al.*, 2017). Per time point, pollen tubes grown from 5 mg of pollen were harvested and freeze-dried overnight. After tissue disruption, 500 µL extraction solution (32.25:12.5:6.25 v/v/v methanol/chloroform/water) were added per sample, vortexed and incubated for 30 minutes at 4 °C under constant shaking. Supernatant was transferred to a new tube and pollen tubes were extracted again with another 500 µL of extraction solution. Incubation was repeated and supernatants combined. 0.0125 mg *allo*-inositol in 0.5 ml water were added, incubated again at 4 °C for 30 min, centrifuged 5 minutes at full speed and the aqueous phase containing the metabolites was transferred. 20 µL were evaporated under N_2_stream and used for derivatisation with 15 µL of methoxyamine hydrochloride in pyridine (30 mg/mL) overnight at room temperature. Derivatisation with 30 µL *N*-methyl-*N*-(trimethylsilyl) trifluoroacetamide (MSTFA) followed for at least 1 h to obtain methoxyimino (MEOX)- and trimethylsilyl (TMS)-derivatives of the metabolites (Bellaire *et al.*, 2014).

For the analysis of sterols, 2 ml of 3:1 v/v MTBE/methanol and 1 ml of 0.9 % w/v NaCl were added to 200 µl of the organic phase and evaporated under N_2_stream. Samples were dissolved in 20 µL pyridine, 10 µl of which were used for MSTFA derivatisation with 10 µL of MSTFA 1-6 h prior to analysis.

### GC-FID and GC-MS

GC-FID analysis of FAMEs was performed as described (Hornung, Pernstich and Feussner, 2002): an Agilent GC 6890 system (Agilent, Waldbronn, Germany) coupled with an FID detector equipped with a capillary DB-23 column (30 m × 0.25 mm, 0.25 µm coating thickness, Agilent, Waldbronn, Germany) was used. Helium served as carrier gas (1 ml/min), with an injector temperature of 220 °C. The temperature gradient was 150 °C for 1 min, 150 – 200 °C at 15 °C min^−1^, 200–250 °C at 2 °C min^−1^, and 250 °C for 10 min.

For quantification, peak integrals were determined using Agilent ChemStation for LC 3D systems (Rev. B.04.03) and used to calculate absolute amounts of TG as well as relative fatty acid contributions.

For the measurement of central metabolites and sterols, GC-MS measurements were performed as previously described in Touraine *et al.* 2019 for metabolites and Berghoff *et al.* 2021 for sterols. If the metabolites were not identified by an external standard, the spectra were identified with the Golm metabolome database (GMD) and the National Institute of Standards and Technology (NIST) spectral library 2.0f. The chemical information on metabolites identified with the GMD can be obtained at http://gmd.mpimp-golm.mpg.de/search.aspx (Kopka *et al.*, 2005). Due to the high levels, sucrose was measured in separate runs using only one-tenth of the sample as used for the regular runs.

Data was analysed using MSD ChemStation (F.01.03.2357). Masses used for quantification are depicted in Supplemental Dataset S7.

### RNA extraction, library preparation, and sequencing

Total RNA was extracted from 200 mg of pollen tubes using Spectrum^TM^ Plant Total RNA Kit (Sigma-Aldrich, St. Louis, MO, USA). RNA-seq libraries were performed using the non-stranded mRNA Kit from Illumina (San Diego, CA, USA; Cat. N°RS-122-2001). Quality and integrity of RNA was assessed with the Fragment Analyzer from Advanced Analytical (Heidelberg, Germany) by using the standard sensitivity RNA Analysis Kit (Agilent, Santa Clara, CA, USA; DNF-471). All samples selected for sequencing exhibited an RNA integrity number over 8. After library generation, for accurate quantitation of cDNA libraries, the fluorometric based system QuantiFluor™dsDNA (Promega, Madison, WI, USA) was used. The size of final cDNA libraries was determined using the dsDNA 905 Reagent Kit (Fragment Analyzer, Advanced Bioanalytical) exhibiting a sizing of 300 bp in average. Libraries were pooled and sequenced on an Illumina HiSeq 4000 (Illumina), generating 50 bp single-end reads (30-40 Mio reads/sample).

### Raw read and quality check

Sequence images were transformed with Illumina software BaseCaller to BCL files, which was demultiplexed to fastq files using bcl2fastq 2.17.1.14. The sequencing quality was asserted using FastQC (version 0.11.5; https://www.bioinformatics.babraham.ac.uk/projects/fastqc/).

### Mapping and normalisation

Samples were aligned to the reference genome *Nicotiana tabacum* (Ntab version TN90, https://www.ncbi.nlm.nih.gov/assembly/GCF_000715135.1/) using the STAR aligner (version 2.5.2a) (Dobin *et al.*, 2013) allowing for 2 mismatches within 50 bases. Subsequently, reads were quantified for all Ntab version TN90 genes in each sample using featureCounts (version 1.5.0-p1) (Liao, Smyth and Shi, 2014).

### Differential gene expression analysis

Read counts were analysed in the R/Bioconductor environment (Release 3.10, www.bioconductor.org) using edgeR package (version 3.28.1) (Robinson, McCarthy and Smyth, 2009; McCarthy, Chen and Smyth, 2012) gene names were translated to UniProt protein identifiers. For more detailed functional analyses, these tobacco proteins were blasted against the TAIR 10 Arabidopsis protein library (TAIR10, https://www.arabidopsis.org/download_files/Proteins/TAIR10_protein_lists/TAIR10_p ep_20101214) using Protein-Protein BLAST 2.5.0+ with a maximum target sequence of 1. Only those hits with an Expect value (E-value, describes amount of hits to be expected by chance for the respective database size) < 10^-5^ were considered. The obtained Arabidopsis AGI-codes were then assigned GO-terms for GO-term analysis The GO term annotations were obtained from the The Arabidopsis Information Resource (www.arabidopsis.org) in a version updated on 1.1.2020.

To analyse genes with a putative involvement in lipid metabolism, a compiled list of lipid genes was generated using the The Arabidopsis Acyl-Lipid Metabolism Website (Li-Beisson *et al.*, 2013), KEGG pathway (https://www.genome.jp/kegg/pathway.html, latest update 10th March 2020), and genes from Kretzschmar *et al.*, 2020, Kelly and Feussner, 2016 and Luttgeharm, Kimberlin and Cahoon, 2016.

### Statistical analyses

Statistical analyses were performed as indicated for the respective experiments. For multiple comparisons, ANOVA was performed, followed by Post-hoc Tukey analysis.

Results are presented as compact letter display of all pair-wise comparisons. Unpaired two-sample t-tests were performed if just two means were compared. Results are presented as * (p < 0.05), ** (p < 0.01) and *** (p < 0.005).

## Supplemental data

Suppl. Figure S1. Micrographs of pollen tube growth under different temperature regimes.

Suppl. Table S1. Parameters for lipid analysis by UPLC-nanoESI-MS/MS. Suppl. Dataset S1. Glycerolipids: Absolute peak areas.

Suppl. Dataset S2. Glycerolipids: Icf-corrected peak areas (partially summed) on the molecular species level and the lipid subclass level.

Suppl. Dataset S3. Glycerolipids: Relative peak areas as % of all icf-corrected peak areas of the respective lipid subclass.

Suppl. Dataset S4. Sphingolipids: Absolute peak areas.

Suppl. Dataset S5. Sphingolipids: Icf-corrected peak areas (partially summed) on the molecular species level and the lipid subclass level.

Suppl. Dataset S6. Sphingolipids: Relative peak areas as % of all icf-corrected peak areas of the respective lipid subclass.

Suppl. Dataset S7. Free sterols: Raw data. Suppl. Dataset S8. Free sterols: Relative levels.

Suppl. Dataset S9. Sterol conjugates: Absolute peak areas.

Suppl. Dataset S10. Sterol conjugates: Icf-corrected peak areas on the molecular species level and the lipid subclass level.

Suppl. Dataset S11. Sterol conjugates: Relative peak areas as % of all icf-corrected peak areas of the respective lipid subclass.

Suppl. Dataset S12. Metabolite analysis: List of analytes.

Suppl. Dataset S13. Metabolite analysis: List of metabolites.

Suppl. Dataset S14. Transcriptomics: Raw Counts.

Suppl. Dataset S15. Transcriptomics: Differential Gene Expression Analysis 3 h 23 °C vs. 6 h 23 °C.

Suppl. Dataset S16. Transcriptomics: Differential Gene Expression Analysis 6 h 23 °C vs. 3 h 23 °C + 3 h 37 °C.

Suppl. Dataset S17. Transcriptomics: GO Term analysis.

Suppl. Dataset S18. Transcriptomics: GO Term analysis - Biological Process.

Suppl. Dataset S19. Transcriptomics: GO Term analysis - Molecular Function.

Suppl. Dataset S20. Transcriptomics: GO Term analysis - Cellular Component.

Suppl. Dataset S21. List of Arabidopsis transcription regulators.

Suppl. Dataset S22. Transcriptomics: List of all detected putative transcription regulators.

Suppl. Dataset S23. List of Arabidopsis genes with a putative involvement in lipid metabolism.

Suppl. Dataset S24. Transcriptomics: List of all detected genes with putative lipid function.

## Acknowledgements

This work was supported by funding from the German research foundation DFG (IS 273/2-2 to T.I., INST 186/1167-1 to Prof. Ivo Feussner), the Studienstiftung des deutschen Volkes (stipends to A.H.R. and P.S.), the IMPRS MolBio (to A.H.R.) and the University of Göttingen by a stipend through the programme “Creativity and Studies” to A.H.R. We are grateful to Sabine Freitag for helping with lipid analysis preparations in the lab.

## References

Allen, J. W. et al. (2018) ‘Induction of oil accumulation by heat stress is metabolically distinct from N stress in the green microalgae *Coccomyxa subellipsoidea* C169’, PLoS ONE, 13(9), pp. 1–20. doi: 10.1371/journal.pone.0204505.

Almeida, J., Perez-Fons, L. and Fraser, P. D. (2021) ‘A transcriptomic, metabolomics and cellular approach to the physiological adaptation of tomato fruit to high temperature’, Plant, Cell & Environment, 44(7), pp. 2211–2229. doi: 10.1111/pce.13854.

Beck, J. G. et al. (2007) ‘Plant sterols in “rafts”: a better way to regulate membrane thermal shocks’, The FASEB Journal, 21(8), pp. 1714–1723. doi: 10.1096/fj.06-7809com.

Bellaire, A. et al. (2014) ‘Metabolism and development - integration of micro computed tomography data and metabolite profiling reveals metabolic reprogramming from floral initiation to silique development’, New Phytologist, 202(1), pp. 322–335. doi: 10.1111/nph.12631.

Berdyshev, E. V. et al. (2005) ‘Quantitative analysis of sphingoid base-1-phosphates as bisacetylated derivatives by liquid chromatography–tandem mass spectrometry’, Analytical Biochemistry, 339(1), pp. 129–136. doi: 10.1016/j.ab.2004.12.006.

Berghoff, S. A. et al. (2021) ‘Microglia facilitate repair of demyelinated lesions via post-squalene sterol synthesis’, Nature Neuroscience, 24(1), pp. 47–60. doi: 10.1038/s41593-020-00757-6.

Boavida, L. C. and McCormick, S. (2007) ‘Temperature as a determinant factor for increased and reproducible *in vitro* pollen germination in *Arabidopsis thaliana*’, Plant Journal, 52(3), pp. 570–582. doi: 10.1111/j.1365-313X.2007.03248.x.

Bonaventure, G. et al. (2003) ‘Disruption of the FATB gene in Arabidopsis demonstrates an essential role of saturated fatty acids in plant growth’, Plant Cell, 15(4), pp. 1020–1033. doi: 10.1105/tpc.008946.

Botte, C. Y. et al. (2011) ‘Chemical inhibitors of monogalactosyldiacylglycerol synthases in *Arabidopsis thaliana*’, Nat Chem Biol, 7(11), pp. 834–842. doi: 10.1038/nchembio.658.

Chen, J. et al. (2006) ‘Characterization of the Arabidopsis thermosensitive mutant *atts02* reveals an important role for galactolipids in thermotolerance’, Plant, Cell and Environment, 29(7), pp. 1437–1448. doi: 10.1111/j.1365-3040.2006.01527.x.

Christenhusz, M. J. M. and Byng, J. W. (2016) ‘The number of known plants species in the world and its annual increase’, Phytotaxa, 261(3), pp. 201–217. doi: 10.11646/phytotaxa.261.3.1.

Coast, O. et al. (2016) ‘Resilience of rice (*Oryza* spp.) pollen germination and tube growth to temperature stress’, Plant Cell and Environment, 39(1), pp. 26–37. doi: 10.1111/pce.12475.

Converti, A. et al. (2009) ‘Effect of temperature and nitrogen concentration on the growth and lipid content of *Nannochloropsis oculata* and *Chlorella vulgaris* for biodiesel production’, Chemical Engineering and Processing: Process Intensification, 48(6), pp. 1146–1151. doi: 10.1016/j.cep.2009.03.006.

Demirel, U. et al. (2020) ‘Physiological, biochemical, and transcriptional responses to single and combined abiotic stress in stress-tolerant and stress-sensitive potato genotypes’, Frontiers in Plant Science, 11(February), pp. 1–21. doi: 10.3389/fpls.2020.00169.

Deruyffelaere, C. et al. (2018) ‘PUX10 is a CDC48A adaptor protein that regulates the extraction of ubiquitinated oleosins from seed lipid droplets in arabidopsis’, Plant Cell, 30(9), pp. 2116–2136. doi: 10.1105/tpc.18.00275.

Ding, P. et al. (2016) ‘Characterization of a pipecolic acid biosynthesis pathway required for systemic acquired resistance’, Plant Cell, 28(10), pp. 2603–2615. doi: 10.1105/tpc.16.00486.

Dobin, A. et al. (2013) ‘STAR: Ultrafast universal RNA-seq aligner’, Bioinformatics, 29(1), pp. 15–21. doi: 10.1093/bioinformatics/bts635.

Dufourc, E. J. (2008) ‘Sterols and membrane dynamics’, Journal of Chemical Biology, 1(1–4), pp. 63–77. doi: 10.1007/s12154-008-0010-6.

Edwards, K. D. et al. (2017) ‘A reference genome for *Nicotiana tabacum* enables map-based cloning of homeologous loci implicated in nitrogen utilization efficiency’, *BMC Genomics*. BMC Genomics, 18(1), pp. 1–14. doi: 10.1186/s12864-017-3791-6.

Flores-Rentería, L. et al. (2018) ‘Higher temperature at lower elevation sites fails to promote acclimation or adaptation to heat stress during pollen germination’, Frontiers in Plant Science, 9(April), pp. 1–14. doi: 10.3389/fpls.2018.00536.

Fragkostefanakis, S. et al. (2016) ‘Unfolded protein response in pollen development and heat stress tolerance’, Plant Reproduction. Springer Berlin Heidelberg, 29(1), pp. 81–91. doi: 10.1007/s00497-016-0276-8.

Ghosh, A. K. et al. (2009) ‘AT4G24160, a soluble acyl-Coenzyme A-dependent lysophosphatidic acid acyltransferase’, Plant Physiology, 151(2), pp. 869–881. doi: 10.1104/pp.109.144261.

Han, B. et al. (2018) ‘Quantitative proteomics and cytology of rice pollen sterol-rich membrane domains reveals pre-established cell polarity cues in mature pollen’, Journal of Proteome Research, 17(4), pp. 1532–1546. doi: 10.1021/acs.jproteome.7b00852.

Haney, C. H. and Long, S. R. (2010) ‘Plant flotillins are required for infection by nitrogen-fixing bacteria’, Proceedings of the National Academy of Sciences of the United States of America, 107(1), pp. 478–483. doi: 10.1073/pnas.0910081107.

Harsh, A. et al. (2016) ‘Effect of short-term heat stress on total sugars, proline and some antioxidant enzymes in moth bean (*Vigna aconitifolia*)’, *Annals of Agricultural Sciences*. Faculty of Agriculture, Ain Shams University, 61(1), pp. 57–64. doi: 10.1016/j.aoas.2016.02.001.

Härtel, H., Dörmann, P. and Benning, C. (2000) ‘DGD1-independent biosynthesis of extraplastidic galactolipids after phosphate deprivation in Arabidopsis’, Proceedings of the National Academy of Sciences of the United States of America, 97(19), pp. 10649–10654. doi: 10.1073/pnas.180320497.

Hedhly, A. (2011) ‘Sensitivity of flowering plant gametophytes to temperature fluctuations’, Environmental and Experimental Botany. Elsevier B.V., 74(1), pp. 9–16. doi: 10.1016/j.envexpbot.2011.03.016.

Henriksen, J. R. et al. (2010) ‘Understanding detergent effects on lipid membranes: A model study of lysolipids’, Biophysical Journal. Biophysical Society, 98(10), pp. 2199–2205. doi: 10.1016/j.bpj.2010.01.037.

Hernández-Vega, J. C. et al. (2017) ‘Detoxification of polycyclic aromatic hydrocarbons (PAHs) in *Arabidopsis thaliana* involves a putative flavonol synthase’, J Hazard Mater, 176(12), pp. 139–148. doi: doi:10.1016/j.jhazmat.2016.08.058.

Herrfurth, C., Liu, Y.-T. and Feussner, I. (2021) ‘Targeted analysis of the plant lipidome by UPLC-NanoESI-MS/MS’, in Bartels D., Dörmann P. (eds) Plant Lipids. Methods in Molecular Biology. Vol 2295. Humana. doi: 10.1007/978-1-0716-1362-7_9.

Higashi, Y. et al. (2015) ‘Landscape of the lipidome and transcriptome under heat stress in *Arabidopsis thaliana*’, Scientific Reports. Nature Publishing Group, 5(1), p. 10533. doi: 10.1038/srep10533.

Higashi, Y. and Saito, K. (2019) ‘Lipidomic studies of membrane glycerolipids in plant leaves under heat stress’, Progress in Lipid Research. Elsevier, 75(March), p. 100990. doi: 10.1016/j.plipres.2019.100990.

Hornung, E., Pernstich, C. and Feussner, I. (2002) ‘Formation of conjugated Δ11 Δ13-double bonds by Δ12-linoleic acid (1,4)-acyl-lipid-desaturase in pomegranate seeds’, European Journal of Biochemistry, 269(19), pp. 4852–4859. doi: 10.1046/j.1432-1033.2002.03184.x.

Ischebeck, T. et al. (2014) ‘Comprehensive cell-specific protein analysis in early and late pollen development from diploid microsporocytes to pollen tube growth’, Molecular & Cellular Proteomics, 13(1), pp. 295–310. doi: 10.1074/mcp.M113.028100.

Ischebeck, T. (2016) ‘Lipids in pollen - They are different’, Biochimica et Biophysica Acta - Molecular and Cell Biology of Lipids. Elsevier B.V., 1861(9), pp. 1315–1328. doi: 10.1016/j.bbalip.2016.03.023.

Ischebeck, T. et al. (2020) ‘Lipid droplets in plants and algae: Distribution, formation, turnover and function’, Seminars in Cell & Developmental Biology, 108(February), pp. 82–93. doi: 10.1016/j.semcdb.2020.02.014.

Jacob, P., Hirt, H. and Bendahmane, A. (2017) ‘The heat-shock protein/chaperone network and multiple stress resistance’, Plant Biotechnology Journal, 15(4), pp. 405– 414. doi: 10.1111/pbi.12659.

James, C. N. et al. (2010) ‘Disruption of the Arabidopsis CGI-58 homologue produces Chanarin-Dorfman-like lipid droplet accumulation in plants’, Proceedings of the National Academy of Sciences of the United States of America, 107(41), pp. 17833– 17838. doi: 10.1073/pnas.0911359107.

Kaplan, F. et al. (2004) ‘Exploring the temperature-stress metabolome’, Plant physiology, 136(December), pp. 4159–4168. doi: 10.1104/pp.104.052142.1.

Karapanos, I. C. et al. (2010) ‘Tomato pollen respiration in relation to *in vitro* germination and pollen tube growth under favourable and stress-inducing temperatures’, Sexual Plant Reproduction, 23(3), pp. 219–224. doi: 10.1007/s00497-009-0132-1.

Keller, M. et al. (2017) ‘Alternative splicing in tomato pollen in response to heat stress’, DNA Research, 24(2), pp. 205–217. doi: 10.1093/dnares/dsw051.

Keller, M. et al. (2018) ‘The coupling of transcriptome and proteome adaptation during development and heat stress response of tomato pollen’, *BMC Genomics*. BMC Genomics, 19(1), pp. 1–20. doi: 10.1186/s12864-018-4824-5.

Kelly, A. a. and Feussner, I. (2016) ‘Oil is on the agenda: Lipid turnover in higher plants’, Biochimica et Biophysica Acta - Molecular and Cell Biology of Lipids. Elsevier B.V., 1861(9), pp. 1253–1268. doi: 10.1016/j.bbalip.2016.04.021.

Kopka, J. et al. (2005) ‘: The Golm metabolome database’, Bioinformatics, 21(8), pp. 1635–1638. doi: 10.1093/bioinformatics/bti236.

Kotak, S. et al. (2007) ‘Complexity of the heat stress response in plants’, Current Opinion in Plant Biology, 10(3), pp. 310–316. doi: https://doi.org/10.1016/j.pbi.2007.04.011.

Kretzschmar, F. K. et al. (2018) ‘PUX10 Is a lipid droplet-localized scaffold protein that interacts with CELL DIVISION CYCLE48 and is involved in the degradation of lipid droplet proteins’, The Plant Cell, 30(9), pp. 2137–2160. doi: 10.1105/tpc.18.00276.

Kretzschmar, F. K. et al. (2020) ‘Identification of low-abundance lipid droplet proteins in seeds and seedlings’, Plant physiology, 182(3), pp. 1326–1345. doi: 10.1104/pp.19.01255.

Lawas, L. M. F. et al. (2019) ‘Metabolic responses of rice cultivars with different tolerance to combined drought and heat stress under field conditions’, GigaScience. Oxford University Press, 8(5), pp. 1–21. doi: 10.1093/gigascience/giz050.

Lee, J. W. et al. (2013) ‘Simultaneous profiling of polar lipids by supercritical fluid chromatography/tandem mass spectrometry with methylation’, Journal of Chromatography A, 1279, pp. 98–107. doi: 10.1016/j.chroma.2013.01.020.

Légeret, B. et al. (2016) ‘Lipidomic and transcriptomic analyses of *Chlamydomonas reinhardtii* under heat stress unveil a direct route for the conversion of membrane lipids into storage lipids’, Plant Cell and Environment, 39(4), pp. 834–847. doi: 10.1111/pce.12656.

Li-Beisson, Y. et al. (2013) ‘Acyl-lipid metabolism.’, The arabidopsis book, 11, p. e0161. doi: 10.1199/tab.0161.

Li, F. et al. (2007) ‘A putative hydroxysteroid dehydrogenase involved in regulating plant growth and development’, Plant Physiology, 145(1), pp. 87–97. doi: 10.1104/pp.107.100560.

Li, R. et al. (2012) ‘A membrane microdomain-associated protein, Arabidopsis Flot1, is involved in a clathrin-independent endocytic pathway and is required for seedling development’, The Plant Cell, 24(5), pp. 2105–2122. doi: 10.1105/tpc.112.095695.

Liao, Y., Smyth, G. K. and Shi, W. (2014) ‘FeatureCounts: An efficient general purpose program for assigning sequence reads to genomic features’, Bioinformatics, 30(7), pp. 923–930. doi: 10.1093/bioinformatics/btt656.

Lu, J. et al. (2020) ‘The role of triacylglycerol in plant stress response’, Plants, 9(4). doi: 10.3390/plants9040472.

Luria, G. et al. (2019) ‘Direct analysis of pollen fitness by flow cytometry: implications for pollen response to stress’, Plant Journal, 98(5), pp. 942–952. doi: 10.1111/tpj.14286.

Luttgeharm, K. D., Kimberlin, A. N. and Cahoon, E. B. (2016) ‘Plant Sphingolipid Metabolism and Function’, in Nakamura, Y. and Li-Beisson, Y. (eds) Lipids in Plant and Algae Development. 86th edn. Springer International Publishing, pp. 249–286.

Luttgeharm, K. D. et al. (2015) ‘Sphingolipid metabolism is strikingly different between pollen and leaf in Arabidopsis as revealed by compositional and gene expression profiling’, Phytochemistry. Elsevier Ltd, 115(1), pp. 121–129. doi: 10.1016/j.phytochem.2015.02.019.

Mamode Cassim, A. et al. (2019) ‘Plant lipids: Key players of plasma membrane organization and function’, Progress in Lipid Research, pp. 1–27. doi: 10.1016/j.plipres.2018.11.002.

Markham, J. E. et al. (2006) ‘Separation and identification of major plant sphingolipid classes from leaves’, Journal of Biological Chemistry, 281(32), pp. 22684–22694. doi: 10.1074/jbc.M604050200.

Mascarenshas, J. P. (1993) ‘Molecular mechanisms of pollen tube growth and differentiation’, 5(October), pp. 1303–1314. doi: https://doi.org/10.1105/tpc.5.10.1303.

Matyash, V. et al. (2008) ‘Lipid extraction by methyl-tert-butyl ether for high-throughput lipidomics’, Journal of Lipid Research, 49(5), pp. 1137–1146. doi: 10.1194/jlr.D700041-JLR200.

McCarthy, D. J., Chen, Y. and Smyth, G. K. (2012) ‘Differential expression analysis of multifactor RNA-Seq experiments with respect to biological variation’, Nucleic Acids Research, 40(10), pp. 4288–4297. doi: 10.1093/nar/gks042.

Michaelson, L. V. et al. (2016) ‘Plant sphingolipids: Their importance in cellular organization and adaption’, Biochimica et Biophysica Acta - Molecular and Cell Biology of Lipids. The Authors, 1861(9), pp. 1329–1335. doi: 10.1016/j.bbalip.2016.04.003.

Mittal, D., Madhyastha, D. A. and Grover, A. (2012) ‘Genome-wide transcriptional profiles during temperature and oxidative stress reveal coordinated expression patterns and overlapping regulons in rice’, PLoS ONE, 7(7), p. e40899. doi: 10.1371/journal.pone.0040899.

Moreno-Pérez, A. J. et al. (2012) ‘Reduced expression of FatA thioesterases in Arabidopsis affects the oil content and fatty acid composition of the seeds’, Planta, 235(3), pp. 629–639. doi: 10.1007/s00425-011-1534-5.

Mueller, S. P. et al. (2015) ‘Accumulation of extra-chloroplastic triacylglycerols in Arabidopsis seedlings during heat acclimation’, Journal of Experimental Botany, 66(15), pp. 4517–4526. doi: 10.1093/jxb/erv226.

Mueller, S. P. et al. (2017) ‘Phospholipid:Diacylglycerol acyltransferase-mediated triacylglyerol synthesis augments basal thermotolerance’, Plant Physiology, 175(1), pp. 486–497. doi: 10.1104/pp.17.00861.

Muhlemann, J. K., Younts, T. L. B. and Muday, G. K. (2018) ‘Flavonols control pollen tube growth and integrity by regulating ROS homeostasis during high-temperature stress’, Proceedings of the National Academy of Sciences of the United States of America, 115(47), pp. E11188–E11197. doi: 10.1073/pnas.1811492115.

Müller, A. O. and Ischebeck, T. (2018) ‘Characterization of the enzymatic activity and physiological function of the lipid droplet-associated triacylglycerol lipase AtOBL1’, New Phytologist, 217(3), pp. 1062–1076. doi: 10.1111/nph.14902.

Müller, F. and Rieu, I. (2016) ‘Acclimation to high temperature during pollen development’, Plant Reproduction, 29(1–2), pp. 107–118. doi: 10.1007/s00497-016-0282-x.

Nakamura, Y., Kobayashi, K. and Ohta, H. (2009) ‘Activation of galactolipid biosynthesis in development of pistils and pollen tubes’, Plant Physiology and Biochemistry, 47(6), pp. 535–539. doi: 10.1016/j.plaphy.2008.12.018.

Návarová, H. et al. (2013) ‘Pipecolic acid, an endogenous mediator of defense amplification and priming, is a critical regulator of inducible plant immunity’, Plant Cell, 24(12), pp. 5123–5141. doi: 10.1105/tpc.112.103564.

Niu, Y. and Xiang, Y. (2018) ‘An overview of biomembrane functions in plant responses to high-temperature stress’, Frontiers in Plant Science, 9(July), pp. 1–18. doi: 10.3389/fpls.2018.00915.

Pérez-Rodríguez, P. et al. (2010) ‘PlnTFDB: updated content and new features of the plant transcription factor database’, Nucleic Acids Research, 38(suppl_1), pp. D822– D827. doi: 10.1093/nar/gkp805.

Pidkowich, M. S. et al. (2007) ‘Modulating seed -ketoacyl-acyl carrier protein synthase II level converts the composition of a temperate seed oil to that of a palm-like tropical oil’, Proceedings of the National Academy of Sciences, 104(11), pp. 4742–4747. doi: 10.1073/pnas.0611141104.

Platre, M. P. et al. (2019) ‘Developmental control of plant Rho GTPase nano-organization by the lipid phosphatidylserine’, Science, 364(6435), pp. 57–62. doi: 10.1126/science.aav9959.

Popko, J. et al. (2016) ‘Metabolome analysis reveals betaine lipids as major source for triglyceride formation, and the accumulation of sedoheptulose during nitrogen-starvation of *Phaeodactylum tricornutum*’, PLoS ONE, 11(10), pp. 1–23. doi: 10.1371/journal.pone.0164673.

Priya, M. et al. (2019) ‘Drought and heat stress-related proteins: an update about their functional relevance in imparting stress tolerance in agricultural crops’, Theoretical and Applied Genetics. Springer Berlin Heidelberg, 132(6), pp. 1607– 1638. doi: 10.1007/s00122-019-03331-2.

Puli, M. R. et al. (2016) ‘Stomatal closure induced by phytosphingosine-1-phosphate and sphingosine-1-phosphate depends on nitric oxide and pH of guard cells in *Pisum sativum*’, Planta, 244(4), pp. 831–841. doi: 10.1007/s00425-016-2545-z.

Rahmati Ishka, M. et al. (2018) ‘A comparison of heat-stress transcriptome changes between wild-type Arabidopsis pollen and a heat-sensitive mutant harboring a knockout of cyclic nucleotide-gated cation channel 16 (*cngc16*)’, *BMC Genomics*. BMC Genomics, 19(1), pp. 1–19. doi: 10.1186/s12864-018-4930-4.

Raja, M. M. et al. (2019) ‘Pollen development and function under heat stress: from effects to responses’, Acta Physiologiae Plantarum. Springer Berlin Heidelberg, pp. 1–20. doi: 10.1007/s11738-019-2835-8.

Read, S. M., Clarke, A. E. and Bacic, A. (1993) ‘Stimulation of growth of cultured *Nicotiana tabacum* W38 pollen tubes by poly(ethylene glycol) and Cu(II) salts’, Protoplasma, 177(1–2), pp. 1–14. doi: 10.1007/BF01403393.

Robinson, M. D., McCarthy, D. J. and Smyth, G. K. (2009) ‘edgeR: A Bioconductor package for differential expression analysis of digital gene expression data’, Bioinformatics, 26(1), pp. 139–140. doi: 10.1093/bioinformatics/btp616.

Rotsch, A. H. et al. (2017) ‘Central metabolite and sterol profiling divides tobacco male gametophyte development and pollen tube growth into eight metabolic phases’, Plant Journal, 92(1), pp. 129–146. doi: 10.1111/tpj.13633.

Santiago, J. P. and Sharkey, T. D. (2019) ‘Pollen development at high temperature and role of carbon and nitrogen metabolites’, Plant Cell and Environment, 42(10), pp. 2759–2775. doi: 10.1111/pce.13576.

Scholz, P. et al. (2020) ‘Signalling pinpointed to the tip: The complex regulatory network that allows pollen tube growth’, Plants, 9(9), p. 1098. doi: 10.3390/plants9091098.

Shao, Q. et al. (2019) ‘New insights into the role of seed oil body proteins in metabolism and plant development’, Frontiers in Plant Science, 10(December), pp. 1–14. doi: 10.3389/fpls.2019.01568.

Shi, W. et al. (2018) ‘Pollen germination and *in vivo* fertilization in response to high-temperature during flowering in hybrid and inbred rice’, Plant Cell and Environment, 41(6), pp. 1287–1297. doi: 10.1111/pce.13146.

Shiva, S. et al. (2020) ‘Leaf lipid alterations in response to heat stress of *Arabidopsis thaliana*’, Plants, 9(7), p. 845. doi: 10.3390/plants9070845.

Simon-Plas, F. et al. (2011) ‘An update on plant membrane rafts’, Current Opinion in Plant Biology, 14(6), pp. 642–649. doi: 10.1016/j.pbi.2011.08.003.

Simons, K. and Ikonen, E. (1997) ‘Functional rafts in cell membranes’, Nature, 387(6633), pp. 569–572. doi: 10.1038/42408.

Snider, J. L. et al. (2011) ‘High temperature limits *in vivo* pollen tube growth rates by altering diurnal carbohydrate balance in field-grown *Gossypium hirsutum* pistils’, Journal of Plant Physiology. Elsevier GmbH., 168(11), pp. 1168–1175. doi: 10.1016/j.jplph.2010.12.011.

Snider, J. L., Oosterhuis, D. M. and Kawakami, E. M. (2011) ‘Diurnal pollen tube growth rate is slowed by high temperature in field-grown *Gossypium hirsutum* pistils’, Journal of Plant Physiology. Elsevier GmbH., 168(5), pp. 441–448. doi: 10.1016/j.jplph.2010.08.003.

Song, G. et al. (2015) ‘Anther response to high-temperature stress during development and pollen thermotolerance heterosis as revealed by pollen tube growth and *in vitro* pollen vigor analysis in upland cotton’, Planta, 241(5), pp. 1271–1285. doi: 10.1007/s00425-015-2259-7.

Spicher, L., Glauser, G. and Kessler, F. (2016) ‘Lipid antioxidant and galactolipid remodeling under temperature stress in tomato plants’, Frontiers in Plant Science, 7(FEB2016), pp. 1–12. doi: 10.3389/fpls.2016.00167.

Sprunck, S. (2020) ‘Twice the fun, double the trouble: gamete interactions in flowering plants’, Current Opinion in Plant Biology. Elsevier Ltd, 53, pp. 106–116. doi: 10.1016/j.pbi.2019.11.003.

Staff, I. A. et al. (1989) ‘Ultrastructural analysis of plastids in angiosperm pollen tubes’, Sexual Plant Reproduction, 2(2), pp. 70–76. doi: 10.1007/BF00191993.

Sunoj, V. S. J. et al. (2017) ‘Resilience of pollen and post-flowering response in diverse sorghum genotypes exposed to heat stress under field conditions’, Crop Science, 57(3), pp. 1658–1669. doi: 10.2135/cropsci2016.08.0706.

Touraine, B. et al. (2019) ‘Iron-sulfur protein NFU2 is required for branched-chain amino acid synthesis in Arabidopsis roots’, Journal of Experimental Botany, 70(6), pp. 1875–1889. doi: 10.1093/jxb/erz050.

Villette, C. et al. (2015) ‘Plant sterol diversity in pollen from angiosperms’, Lipids, 50(8), pp. 749–760. doi: 10.1007/s11745-015-4008-x.

Wu Jingrui et al. (1994) ‘A mutant of Arabidopsis deficient in the elongation of palmitic acid’, Plant Physiology, 106(1), pp. 143–150. doi: 10.1104/pp.106.1.143.

Yang, H. et al. (2007) ‘The arabidopsis BAP1 and BAP2 genes are general inhibitors of programmed cell death’, Plant Physiology, 145(1), pp. 135–146. doi: 10.1104/pp.107.100800.

Yang, Y. et al. (2013) ‘At high temperature lipid production in *Ettlia oleoabundans* occurs before nitrate depletion’, Applied Microbiology and Biotechnology, 97(5), pp. 2263–2273. doi: 10.1007/s00253-012-4671-2.

Zhang, C. et al. (2018) ‘Heat stress induces spikelet sterility in rice at anthesis through inhibition of pollen tube elongation interfering with auxin homeostasis in pollinated pistils’, Rice. Rice, 11(1), p. 14. doi: 10.1186/s12284-018-0206-5.

Zhang, Z. et al. (2016) ‘OsHSD1 , a hydroxysteroid dehydrogenase, is involved in cuticle formation and lipid homeostasis in rice’, Plant Science, 249, pp. 35–45. doi: 10.1016/j.plantsci.2016.05.005.

Zhao, Q. et al. (2018) ‘Relationship of ROS accumulation and superoxide dismutase isozymes in developing anther with floret fertility of rice under heat stress’, Plant Physiology and Biochemistry, 122, pp. 90–101. doi: 10.1016/j.plaphy.2017.11.009.

Zinta, G. et al. (2018) ‘Dynamics of metabolic responses to periods of combined heat and drought in *Arabidopsis thaliana* under ambient and elevated atmospheric CO_2_’, Journal of Experimental Botany, 69(8), pp. 2159–2170. doi: 10.1093/jxb/ery055.

